# The disappointment centre of the brain gets exciting: A systematic review of habenula dysfunction in depression

**DOI:** 10.1101/2024.04.15.589608

**Authors:** Sarah Cameron, Katrina Weston-Green, Kelly A Newell

## Abstract

**Background:** The habenula is an epithalamic brain structure that acts as a neuroanatomical hub connecting the limbic forebrain to the major monoamine centres. Abnormal habenula activity is increasingly implicated in depression, with a surge in publications on this topic in the last 5 years. Direct stimulation of the habenula is sufficient to induce a depressive phenotype in rodents, suggesting a causative role in depression. However, the molecular basis of habenula dysfunction in depression remains elusive and it is unclear how the preclinical advancements translate to the clinical field.

**Methods:** A systematic literature search was conducted following the PRISMA guidelines. The two search terms depress* and habenula* were applied across the databases Scopus, Web of Science and PubMed. Studies eligible for inclusion must have examined changes in the habenula in clinical cases of depression or preclinical models of depression.

**Results:** Preclinical studies (n=57) measured markers of habenula activity (n=16) and neuronal firing (n=21), largely implicating habenula hyperactivity in depression. Neurotransmission was briefly explored (n=13), suggesting imbalances within excitatory and inhibitory habenula signalling. Additional preclinical studies reported neuroconnectivity (n=1), inflammatory (n=2), genomic (n=2) and circadian rhythm (n=2) abnormalities. Seven preclinical studies (12.2%) included both males and females. From these, 5 studies (71%) reported a significant difference between the sexes in at least one habenula measure taken. Clinical studies (n=18) reported abnormalities in habenula connectivity (n=11), volume (n=5) and molecular markers (n=2). Clinical studies generally included male and female subjects (n=15), however, few of these studies examined sex as a biological variable (n=5)

**Conclusions:** Both preclinical and clinical evidence suggest the habenula is disrupted in depression. However, there are opportunities for sex-specific analyses across both areas. Preclinical evidence consistently suggests habenula hyperactivity as a primary driver for the development of depressive symptoms. Clinical studies support gross habenula abnormalities such as altered activation, connectivity, and volume, with emerging evidence of blood brain barrier dysfunction, however, progress is limited by a lack of detailed molecular analyses.

## 1.0 Introduction

Depression is a highly prevalent and debilitating mood disorder, affecting over 300 million individuals and is the leading cause of disability worldwide (1). Depression is characterised by a long disease course, high suicide rates and symptoms that include depressed mood, an inability to feel pleasure (anhedonia), sleep disturbances, impaired cognition and lethargy (1,2). Women are disproportionately affected and account for approximately two-thirds of the clinical population (3). Despite the tremendous contribution to the global burden of disease, current pharmacological treatment options for depression are associated with a wide range of side effects (4,5), have a slow onset of action and low efficacy leading to a high incidence of treatment resistance (6,7). Due to the complex and heterogeneous nature of depression, the underlying molecular mechanisms remain unclear. Novel therapeutic targets are needed to reduce the social, economic, and psychological burden associated with the symptoms of depression and provide reprieve for those living with the illness.

Recently an evolutionary conserved, epithalamic brain structure known as the habenula (Hb) has garnered substantial interest in the context of mood disorders, particularly, depression. The past 5 years has seen a surge in publications (***Figure 1***) as researchers seek to understand how the anatomically tiny, yet functionally vital, Hb goes awry in depression. The Hb is comprised of two divisions, the medial (MHb) and lateral Hb (LHb) (8). The LHb, colloquially termed the “anti-reward” or “disappointment” centre of the brain, is involved in the regulation of motivational states (9), stress adaptation (10), sleep (11), and the encoding of negative stimuli (12). Our understanding of the molecular and cellular properties of the Hb is predominately derived from rodent studies (13–15). Primarily comprised of glutamatergic neurons, the LHb receives inputs from the basal ganglia and limbic forebrain and projects onto major monoamine centres including the dopaminergic ventral tegmental area (VTA) and serotonergic dorsal raphe nucleus (DRN) (13). Greater neuronal activity within the LHb leads to an inhibition within each of these regions, making it a key point of convergence modulating monoamine transmission (13). Compared to the LHb, the function of the MHb is relatively unclear and unexplored and in-vivo electrophysiology studies of the MHb are challenged by the small size of the subregion (13). Like the LHb, the MHb is primarily comprised of glutamatergic neurons; however, a large portion of MHb neurons also co-release acetylcholine and the neuropeptide, substance P (16). The MHb projects onto the interpeduncular nucleus (IPN) (13) and has been linked to depression through its downstream regulatory effects on the dopaminergic and serotonergic systems via the IPN-DRN and IPN-VTA circuitry (17). In addition, the MHb is involved in the regulation of fear and anxiety (18) and has been heavily implicated in nicotine addiction (19). The human Hb exhibits similar neuroanatomical connectivity to rodents, however proportionately the LHb is significantly larger than the MHb, comprising up to 95% of the total Hb volume, comparatively, the rodent LHb makes up approximately 60% of the total Hb volume (20, 21). The relative size difference may suggest a more complex modulation within the human LHb, highlighting the importance of clinical research.

**Figure 1.**
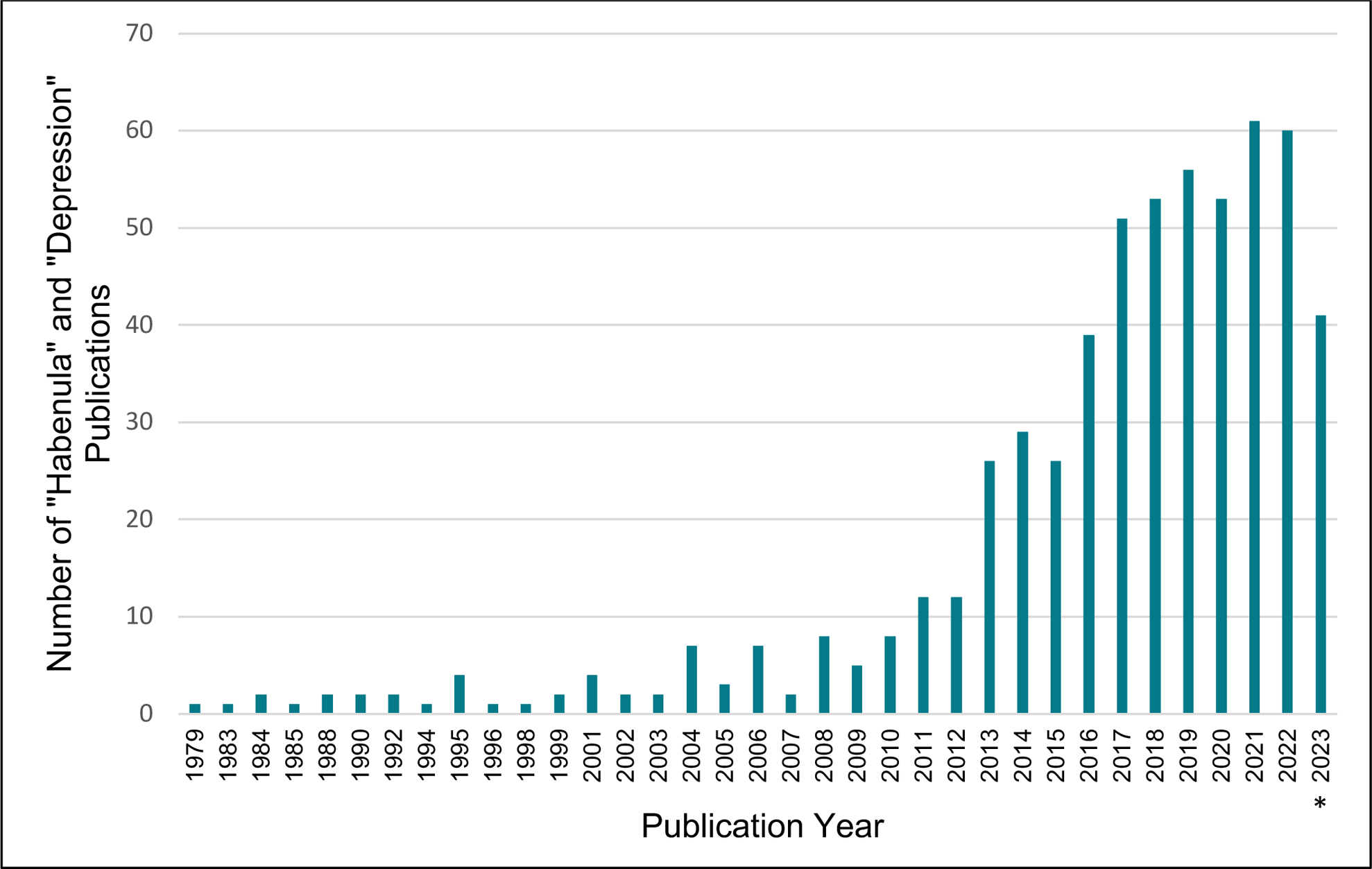
Graph illustrating the number of “habenula” and “depression” publications from 1979 to 2023 taken from PubMed database. *Data taken from Jan-Aug 2023.

The majority of literature reporting on the Hb is centred on preclinical investigations. In rodents, direct stimulation of the LHb is sufficient to induce a depressive-like phenotype (22, 23). Further, inhibition of the LHb via optogenetics and local administration of rapid-acting anti-depressants (ketamine) can reverse these behavioural deficits (24, 25). Emerging research has identified sexual dimorphisms in the Hb circuitry (26, 27). Specifically, female mice were reported to exhibit stronger excitatory inputs; conversely, males displayed greater GABAergic afferent projections to the LHb (27). Despite this, the literature exploring whether there are sex differences in the Hb in clinical depression or preclinical models of depression is sparse.

Despite the recent surge in interest in the Hb, the molecular, cellular and circuit properties of this region remain elusive in humans and it is unclear if the remarkable developments observed in preclinical models translates to the human brain. Additionally, it’s important to highlight a variety of approaches are often used to induce depressive-like behaviours in preclinical models; these range from environmental insults to selective breeding and genetic manipulation (29). As a result, the question on whether Hb dysfunction is consistently observed across each of these models remains unanswered. The current systematic review analyses and evaluates where preclinical and clinical fields support and oppose one another and discusses the current state of Hb research in the context of depression, as well as considerations moving towards future therapeutics.

## 2.0 Methods

The systematic literature review was conducted adhering to the Preferred Reporting Items for Systematic Reviews and Meta-Analyses (PRISMA) 2020 guidelines (Supp Table 1). Articles retained and excluded were recorded in a four-phase flow diagram following the PRISMA statement requirements (***Figure 2***) (30).

**Figure 2.**
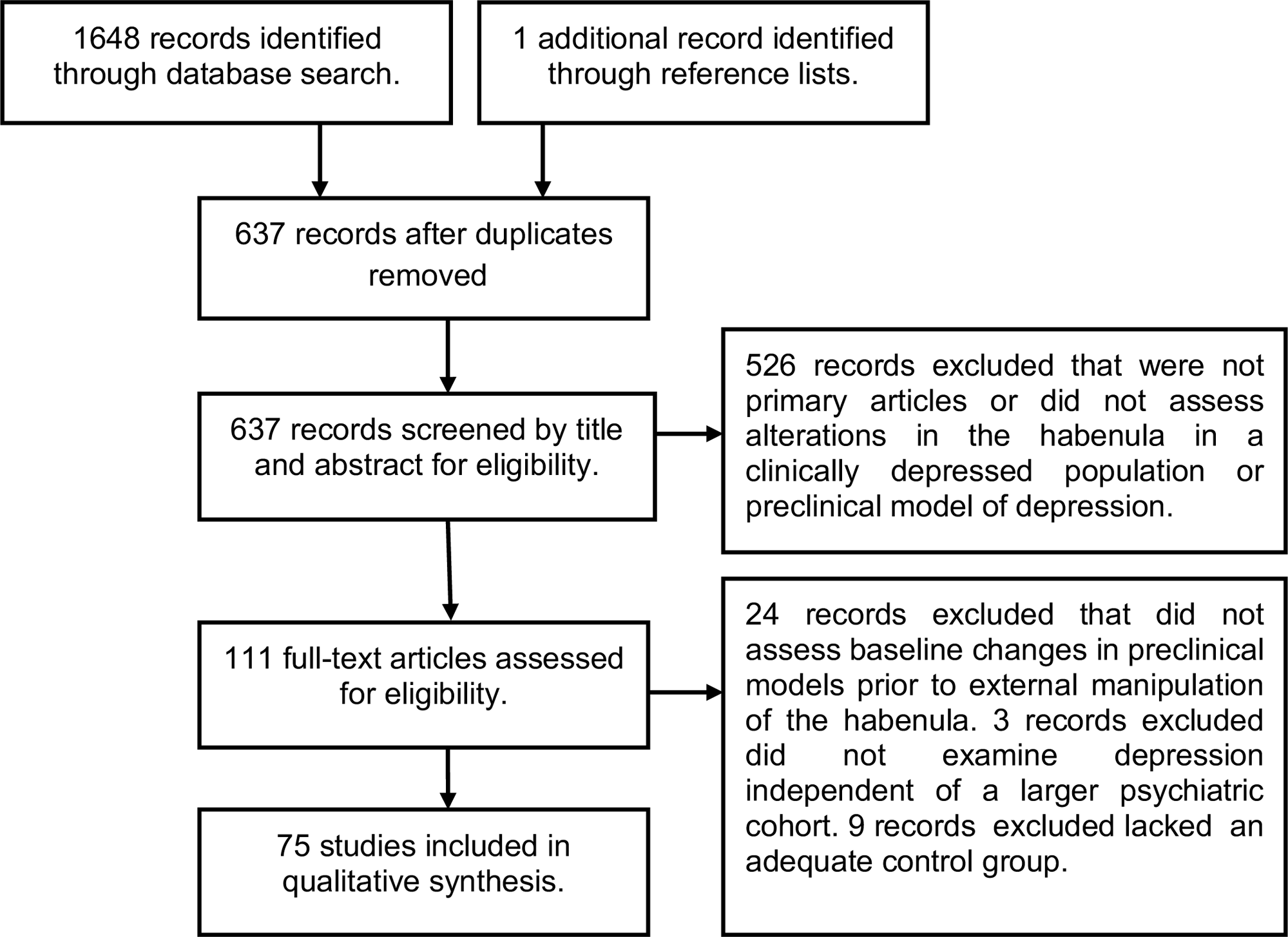
PRISMA Flow Diagram identifying the studies meeting the inclusion criteria for systematic review.

**Table 1.**
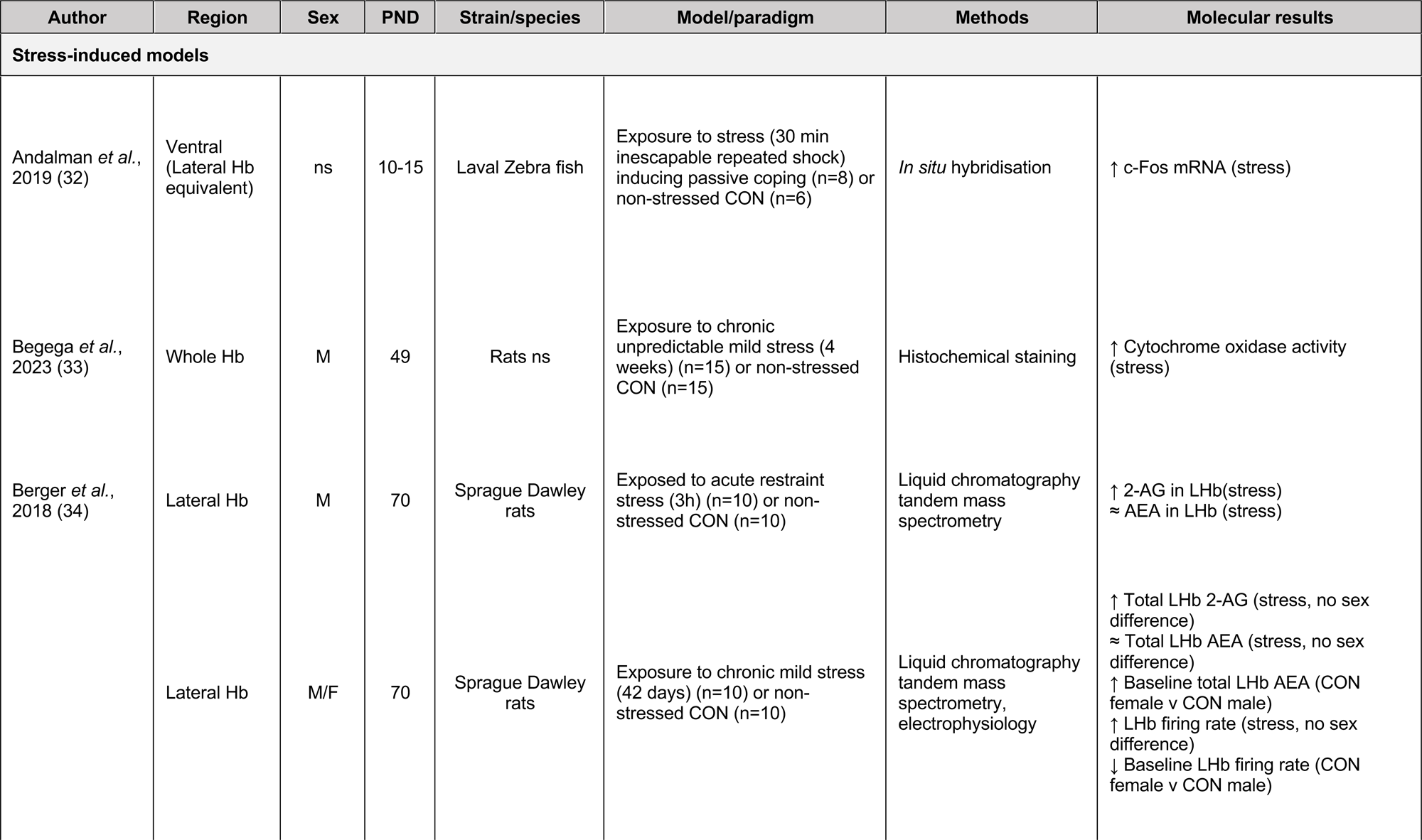
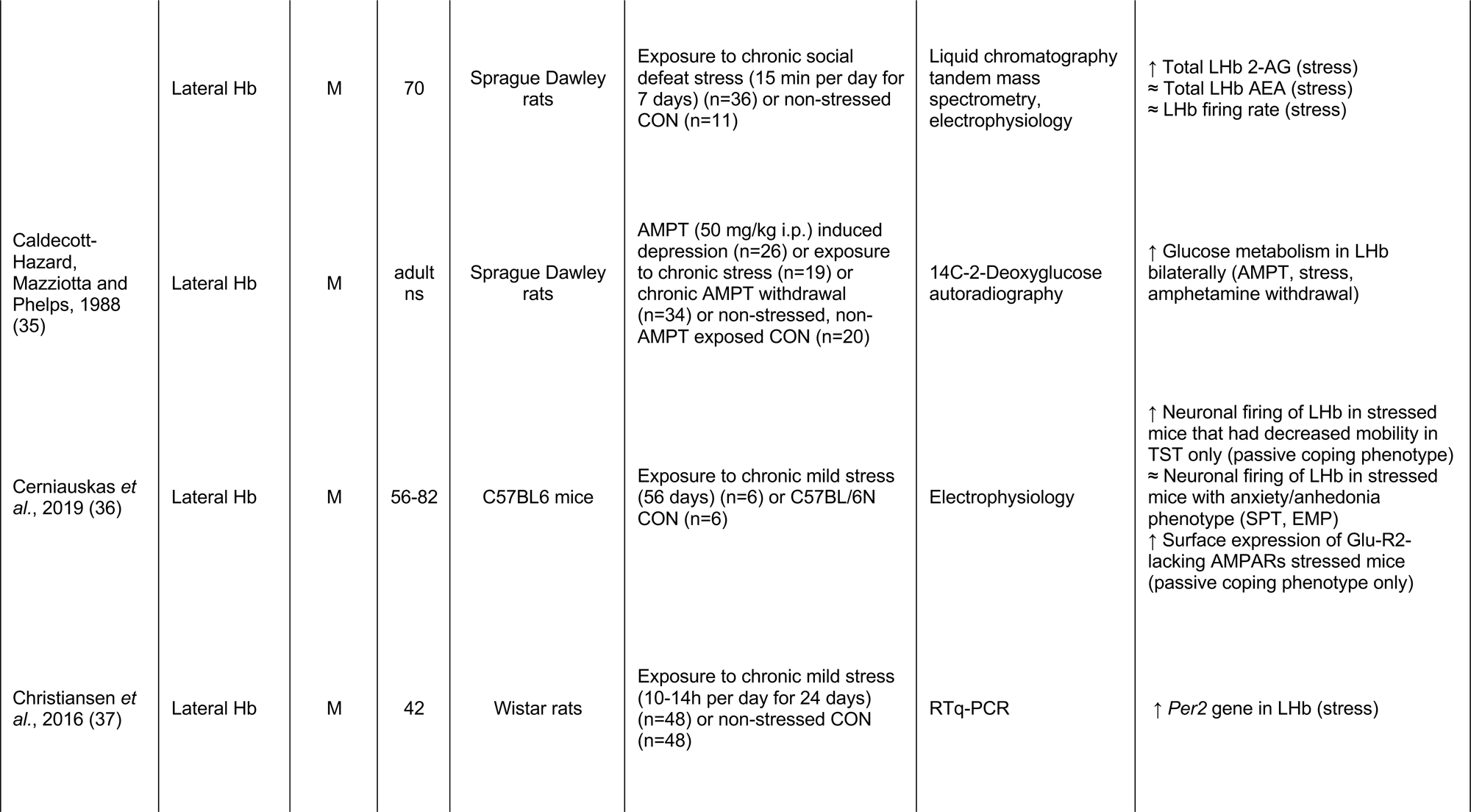
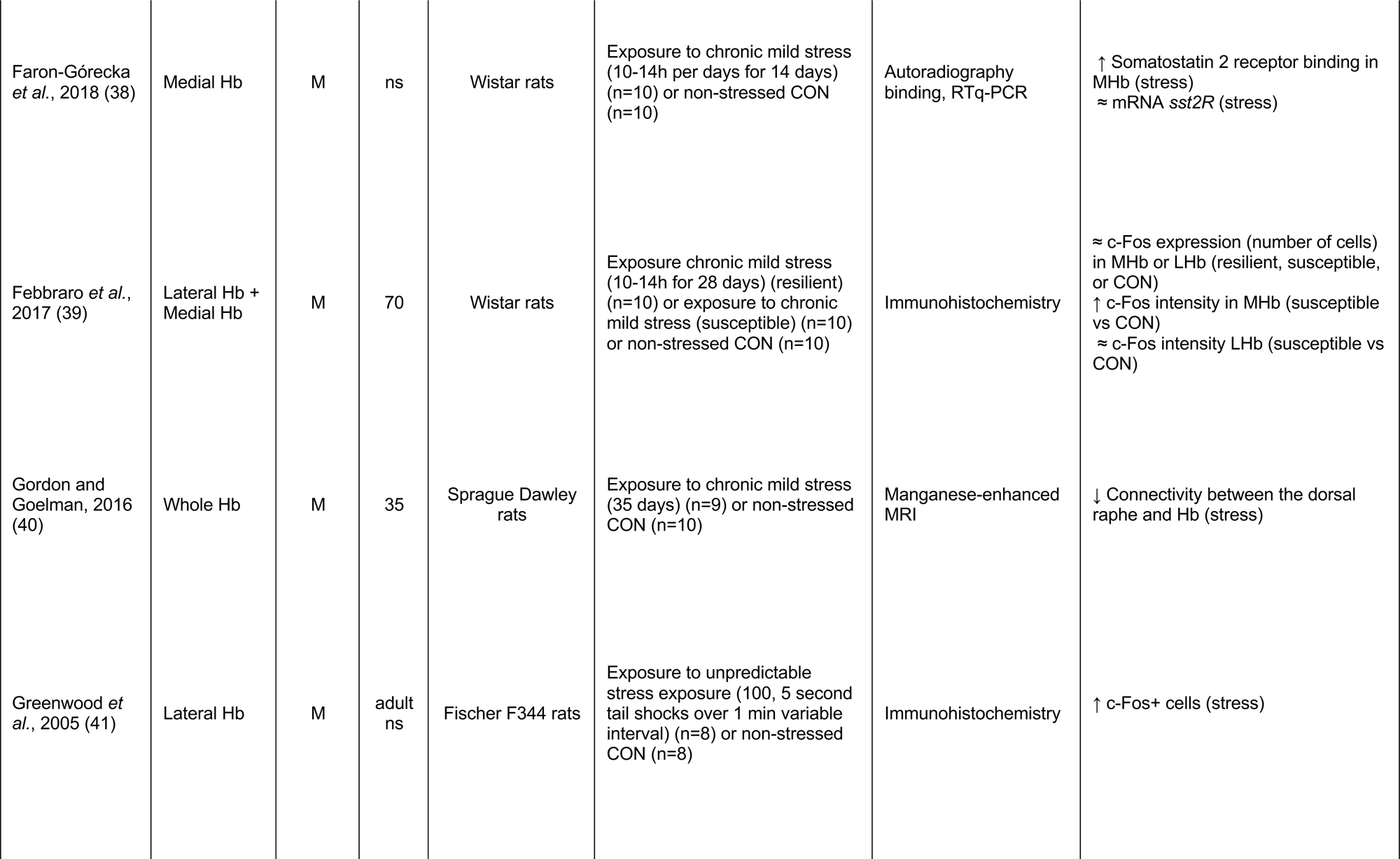
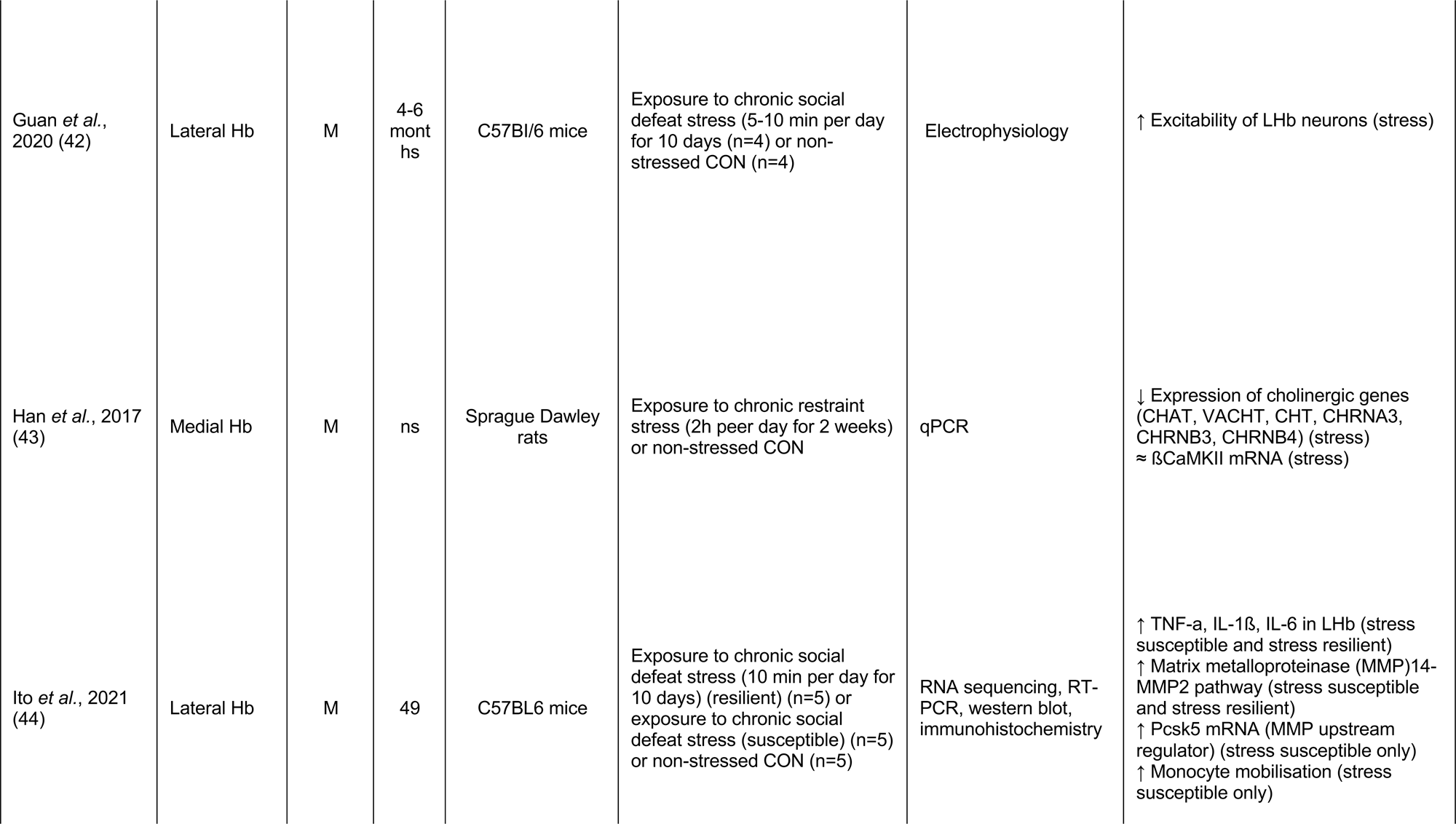
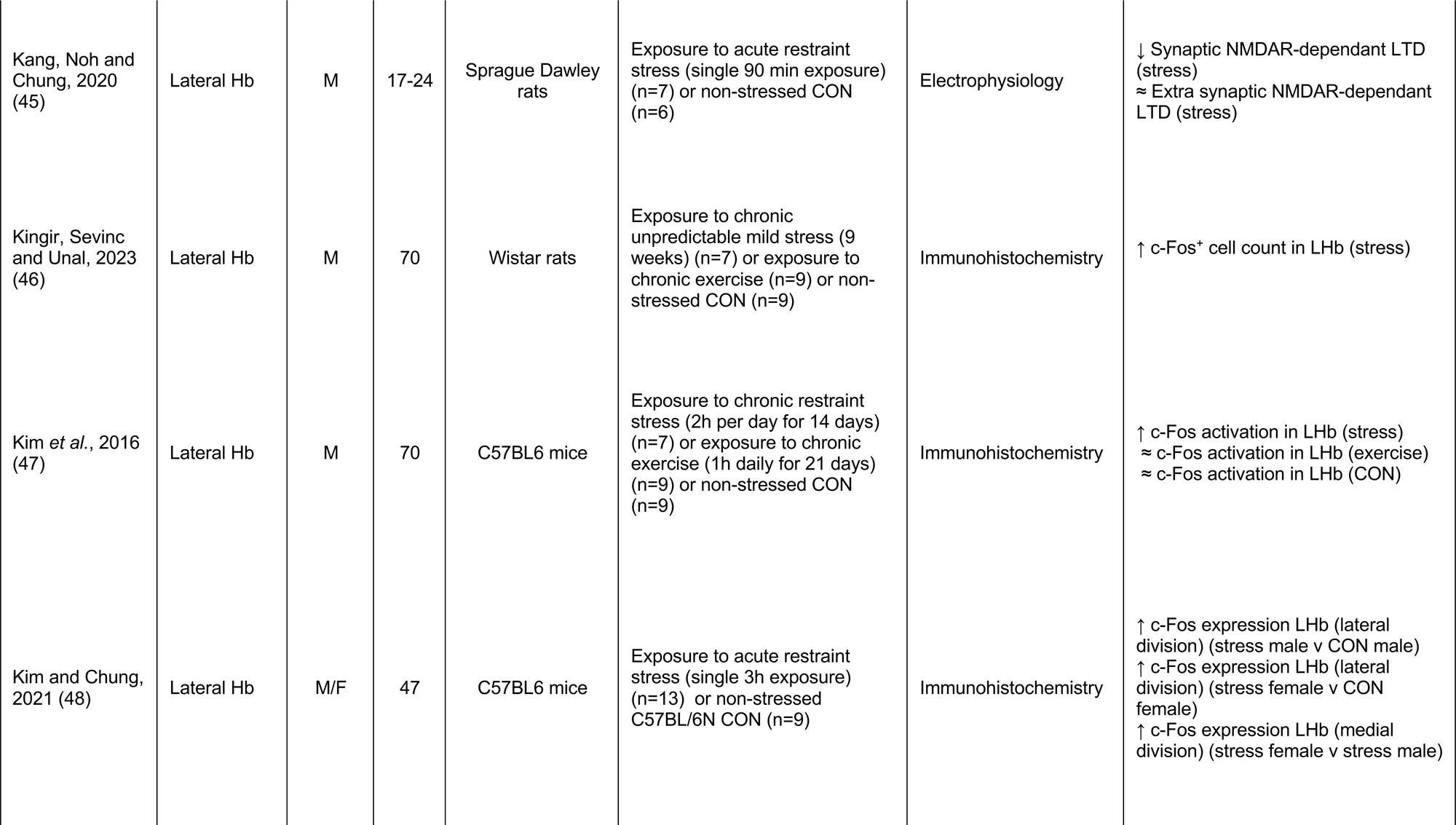
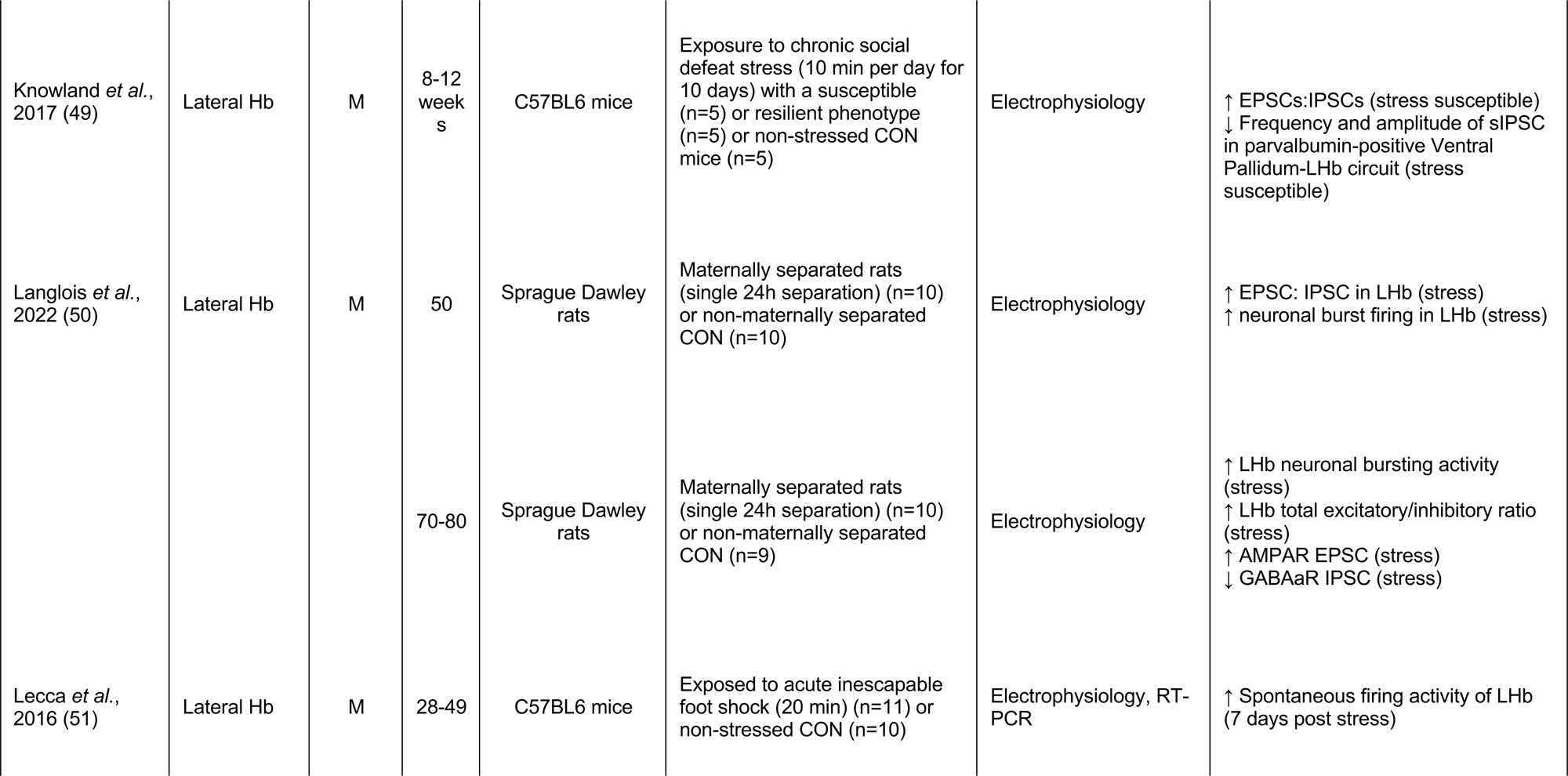
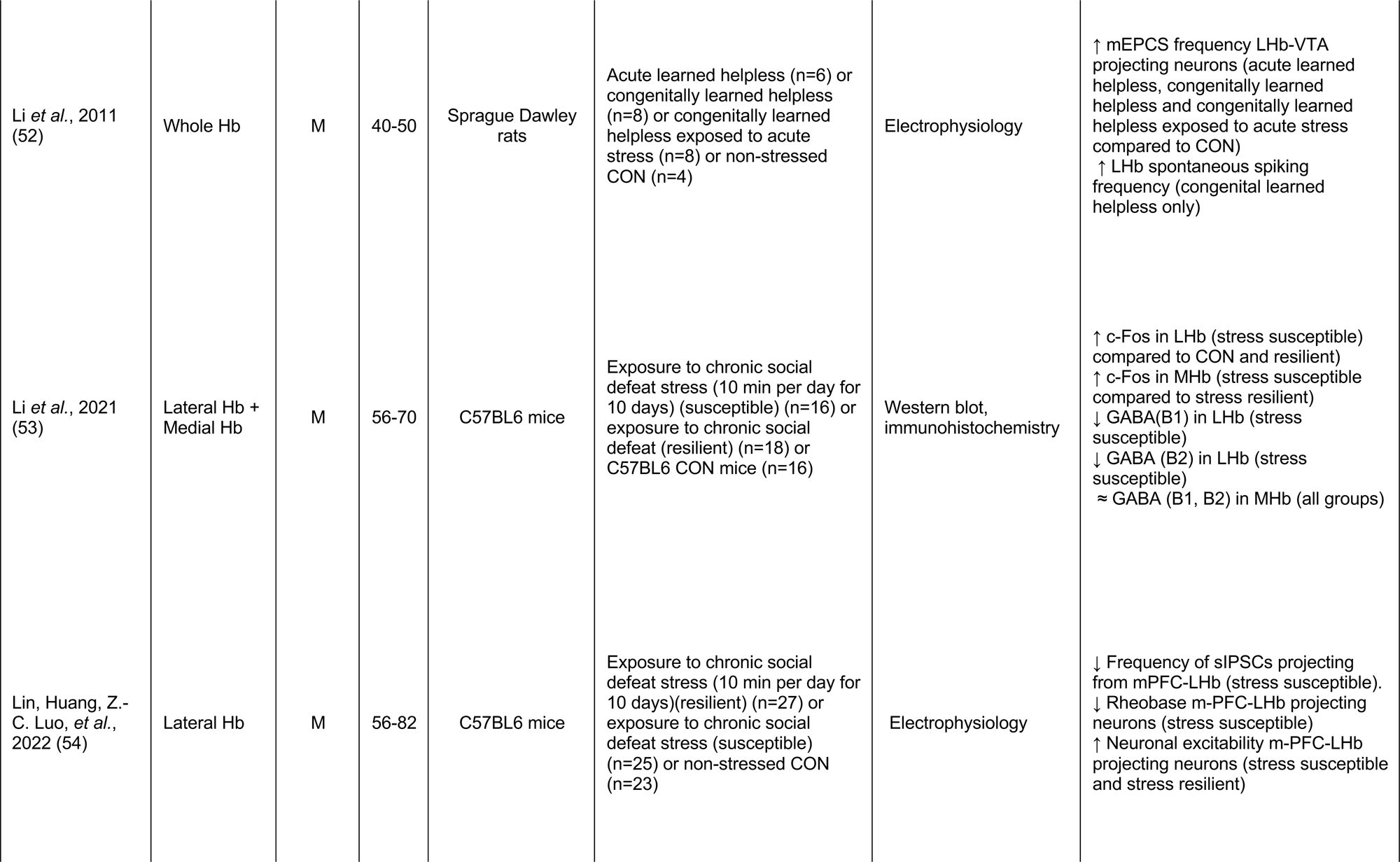
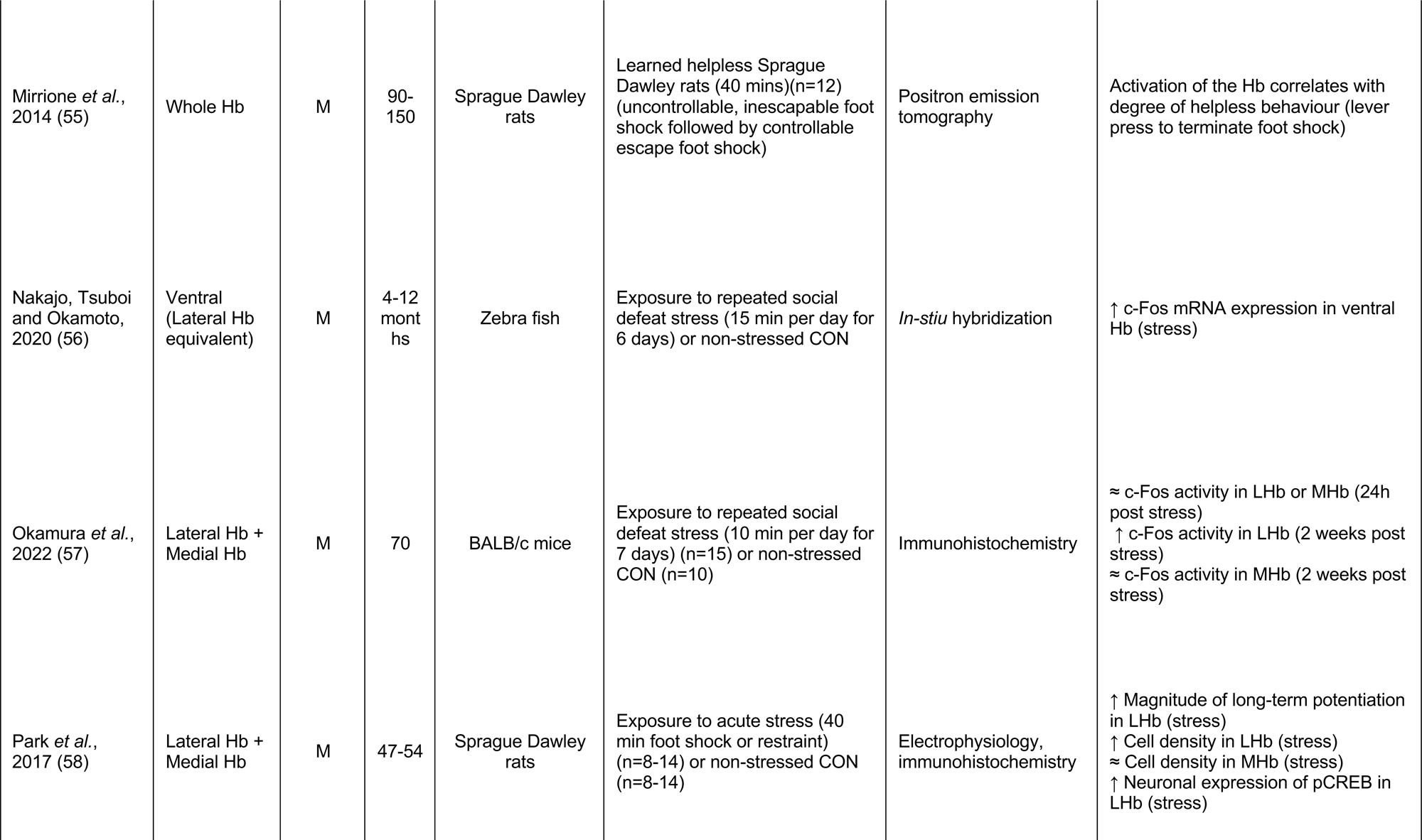
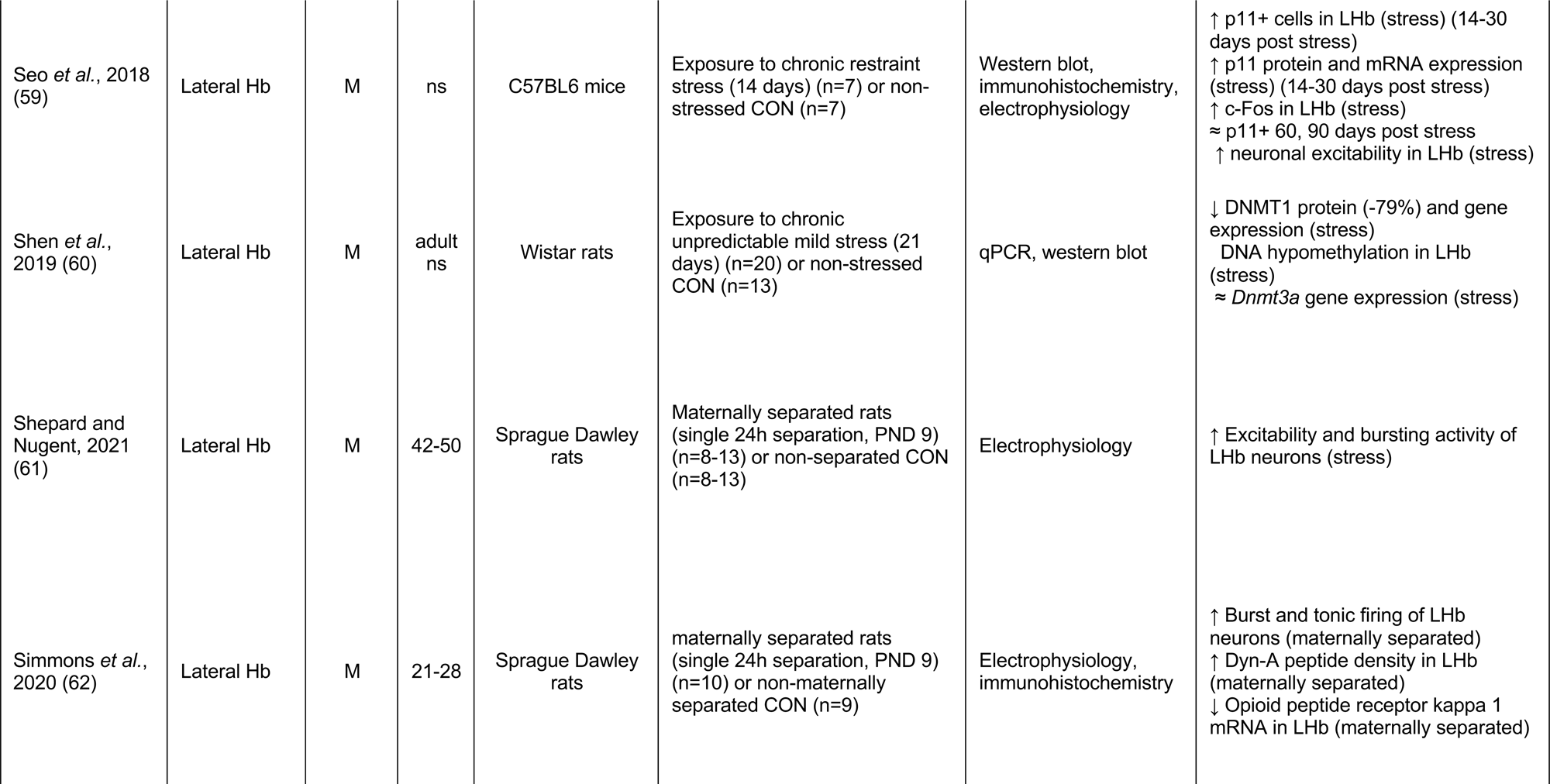
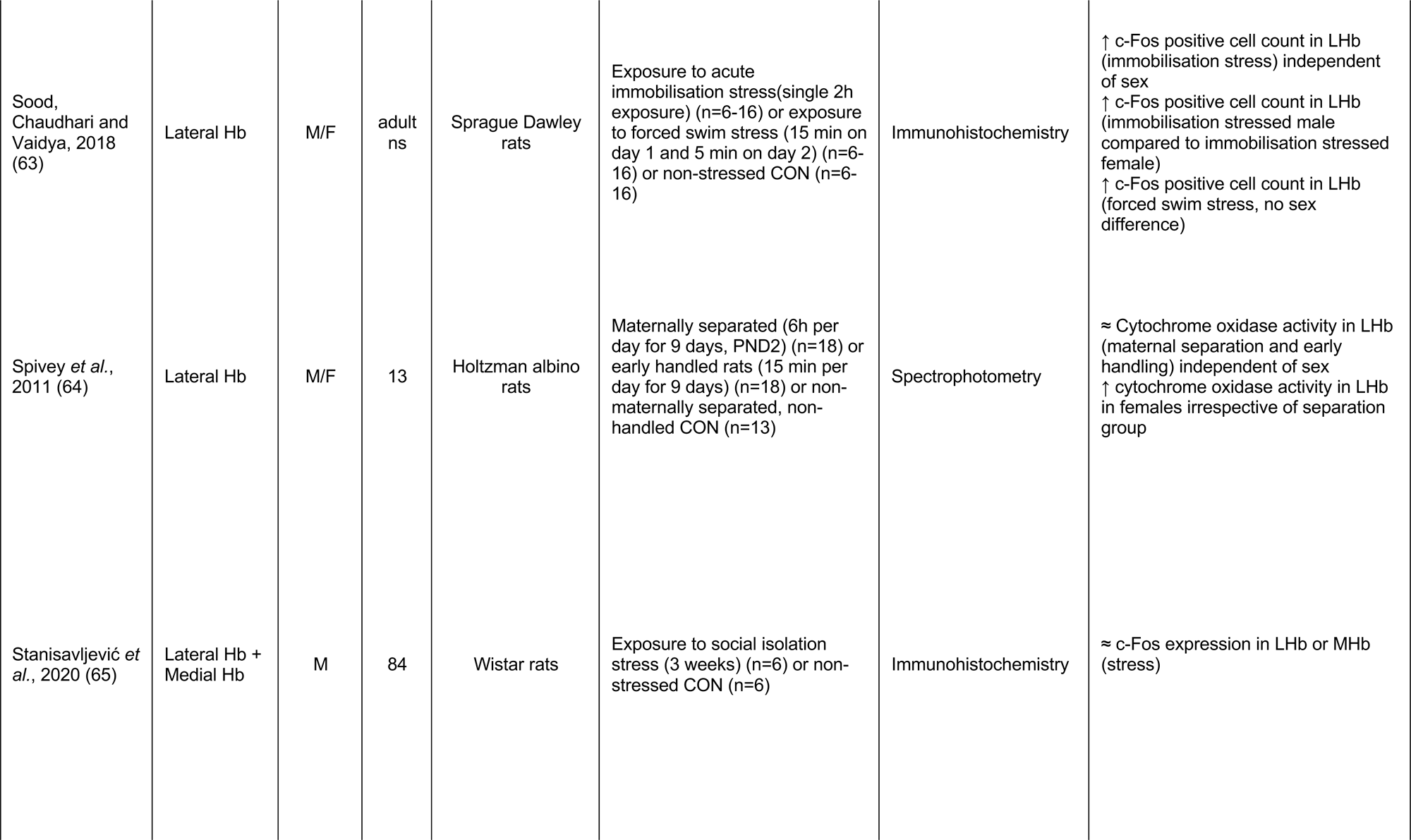
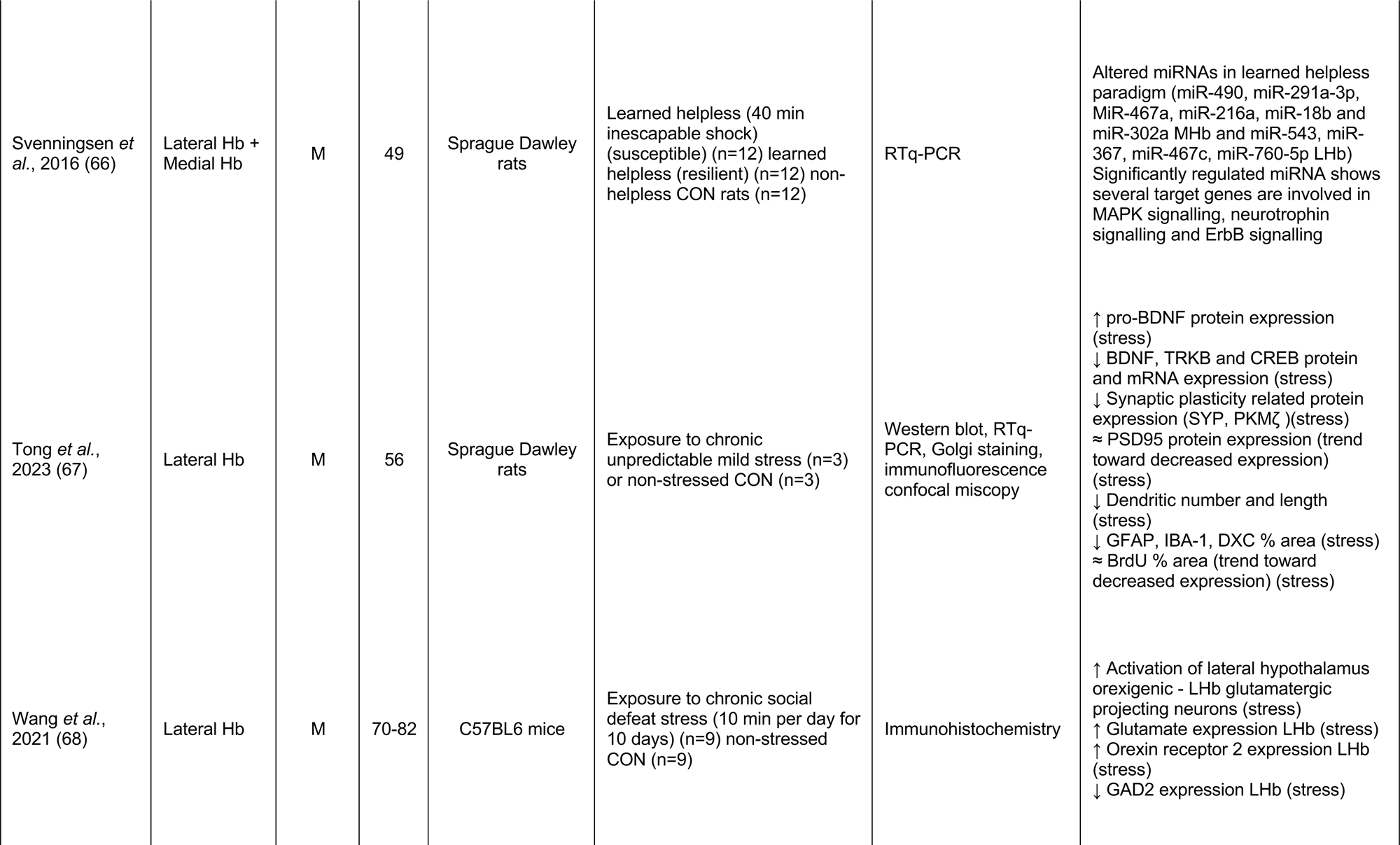
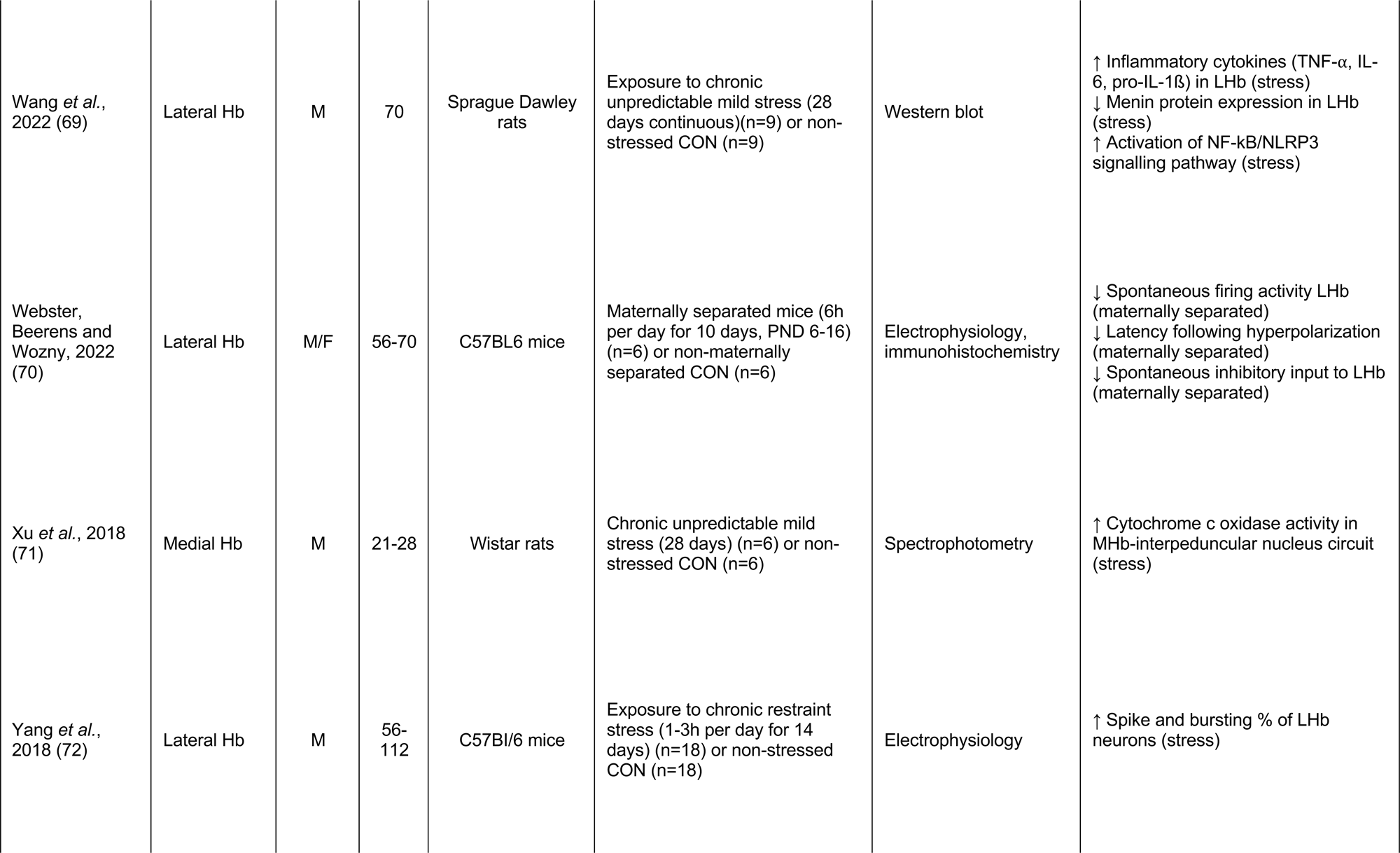
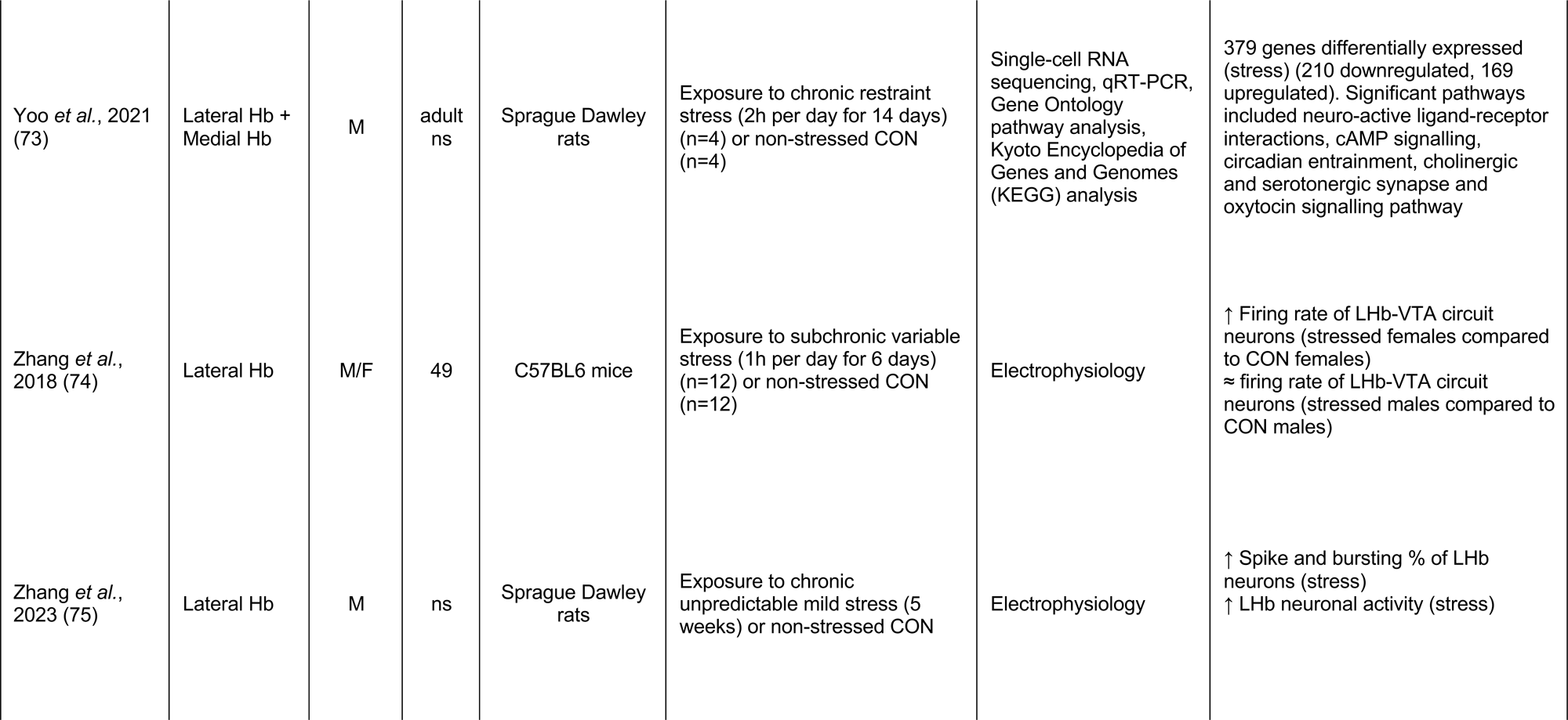
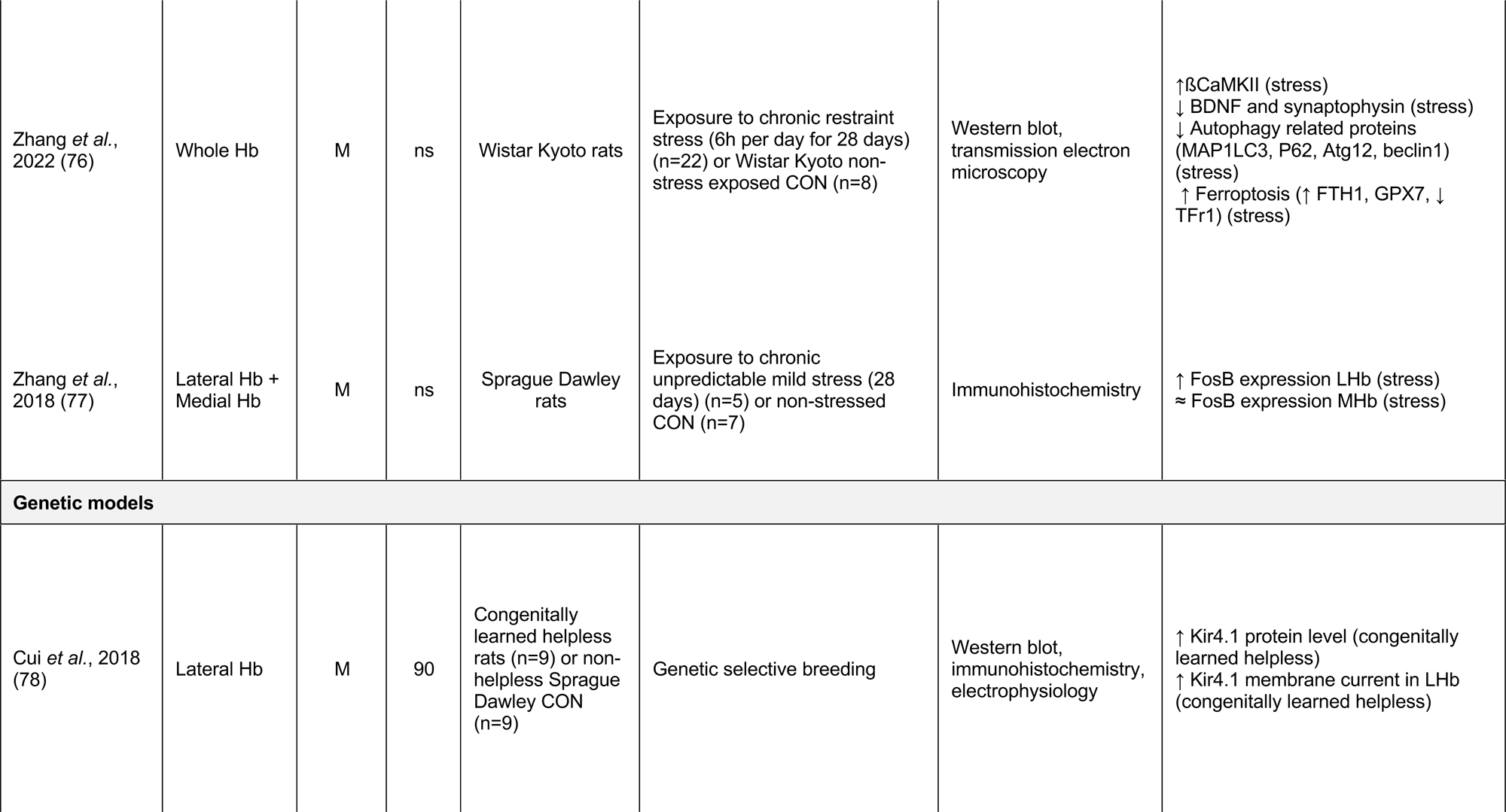
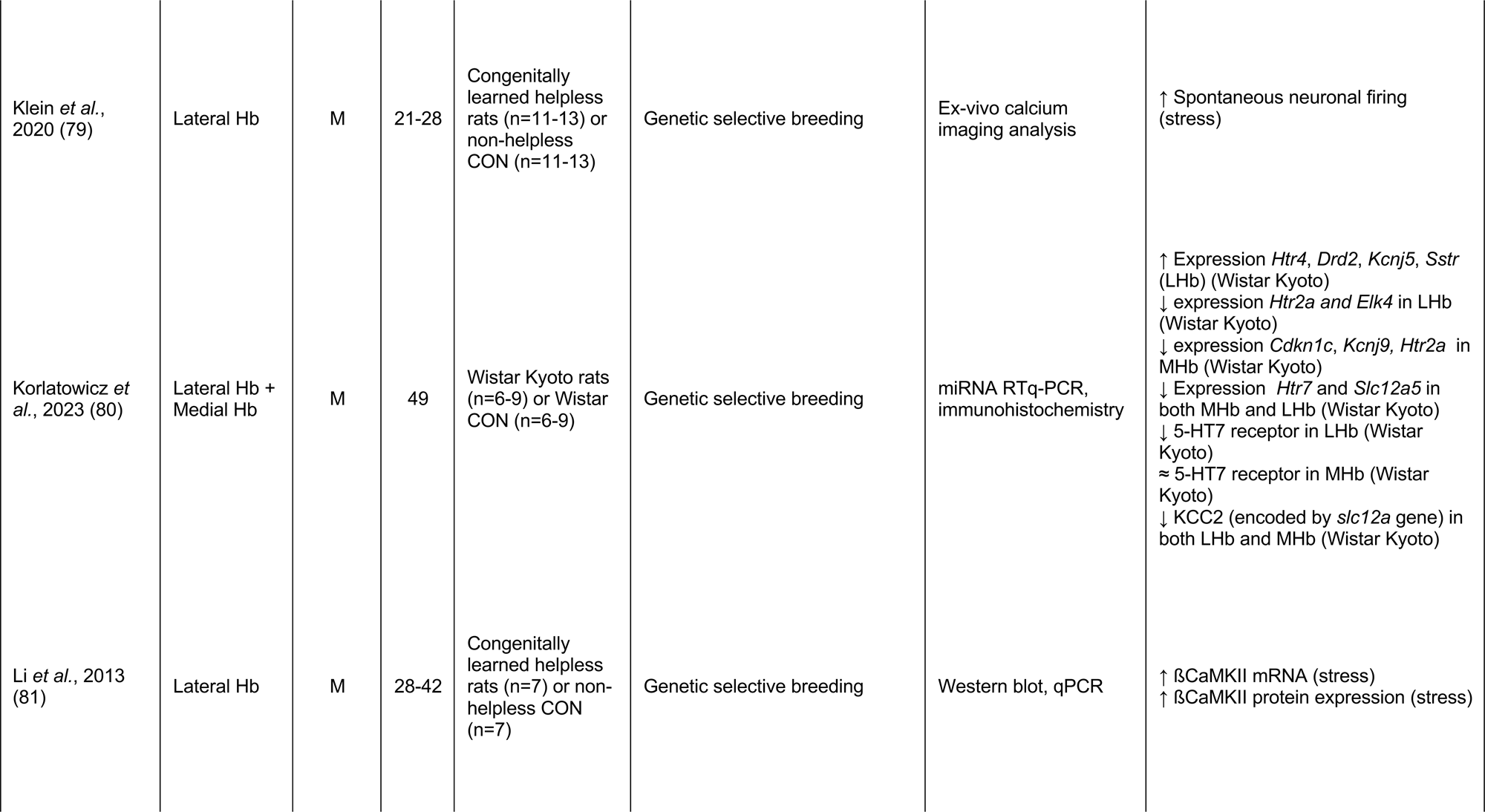
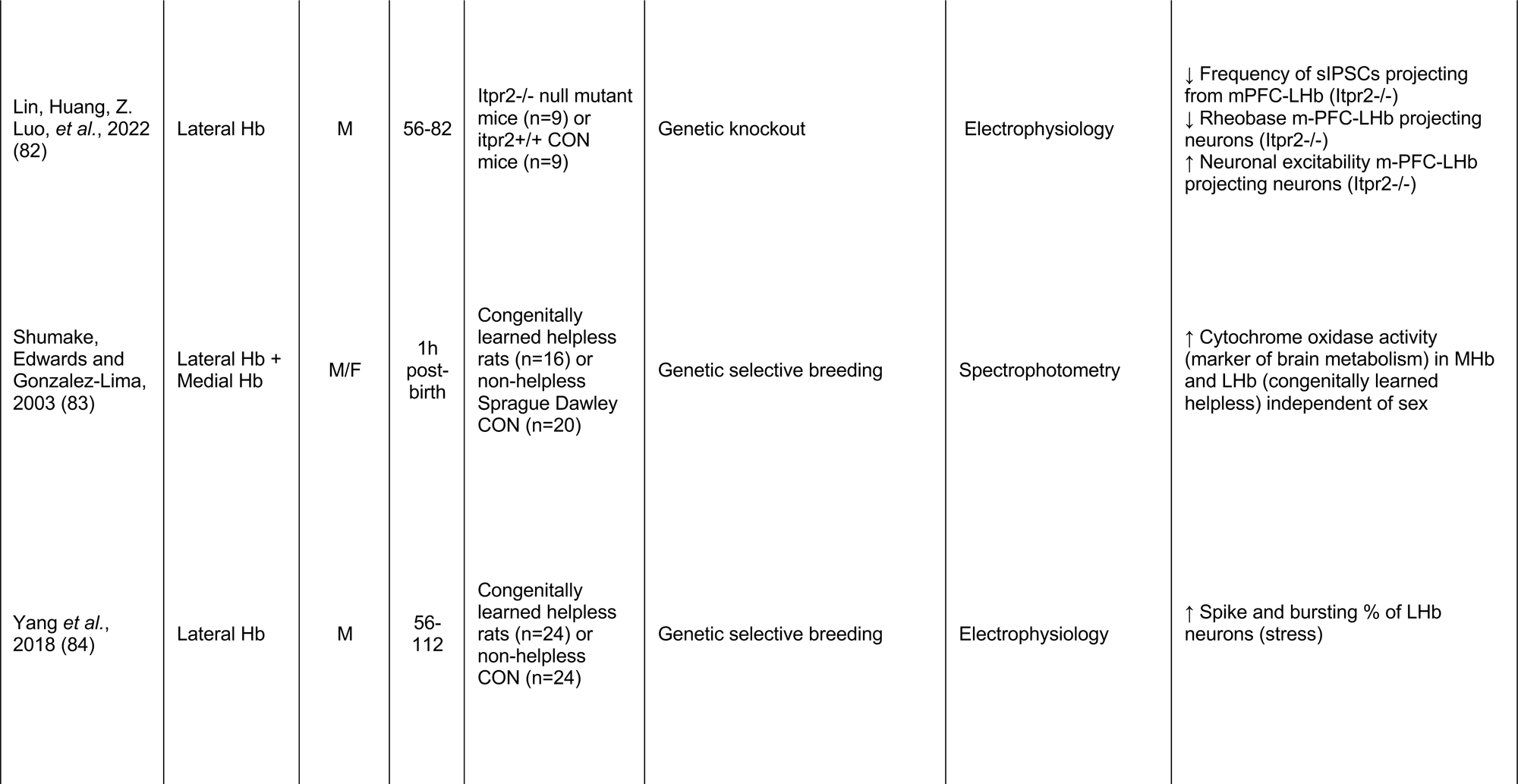
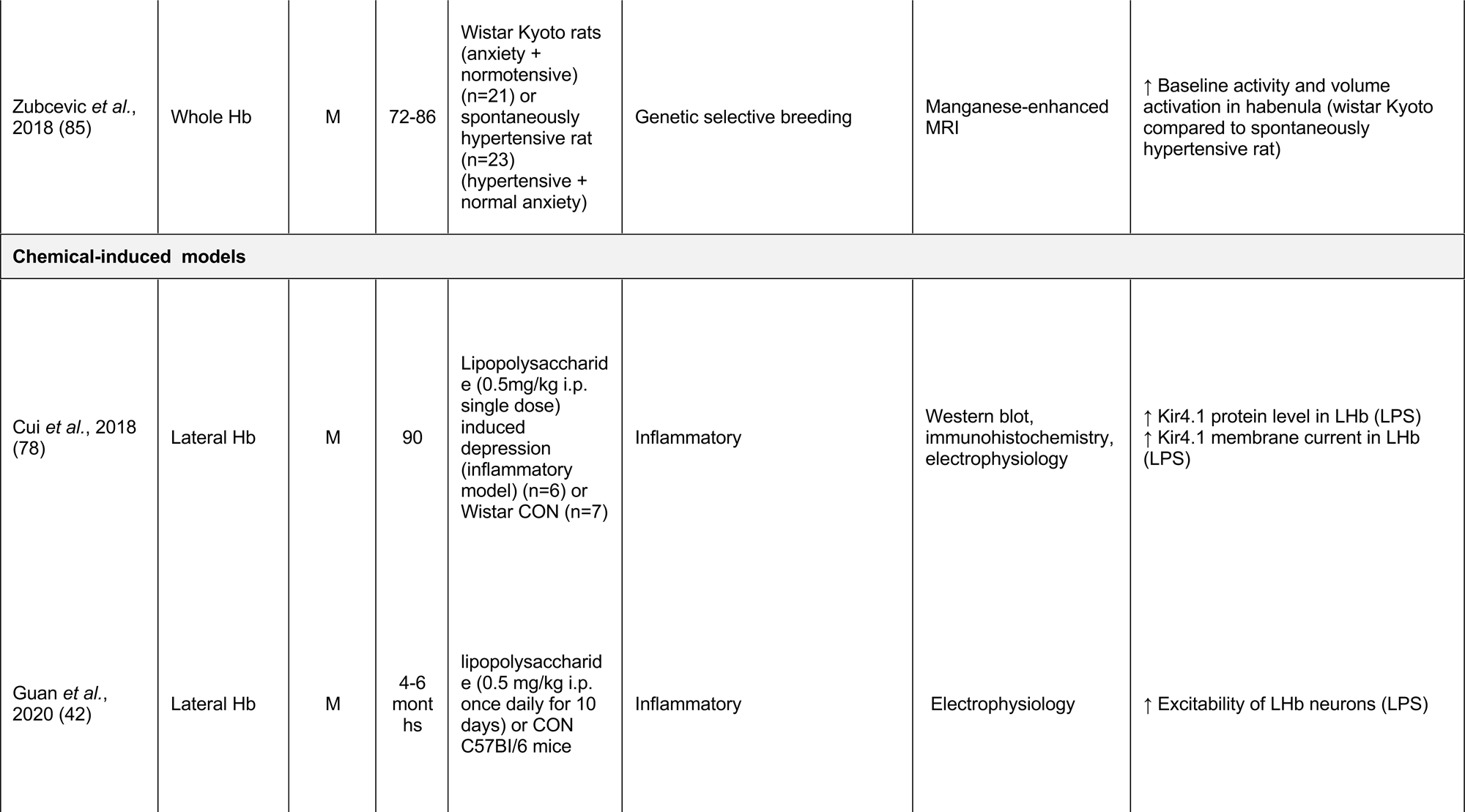
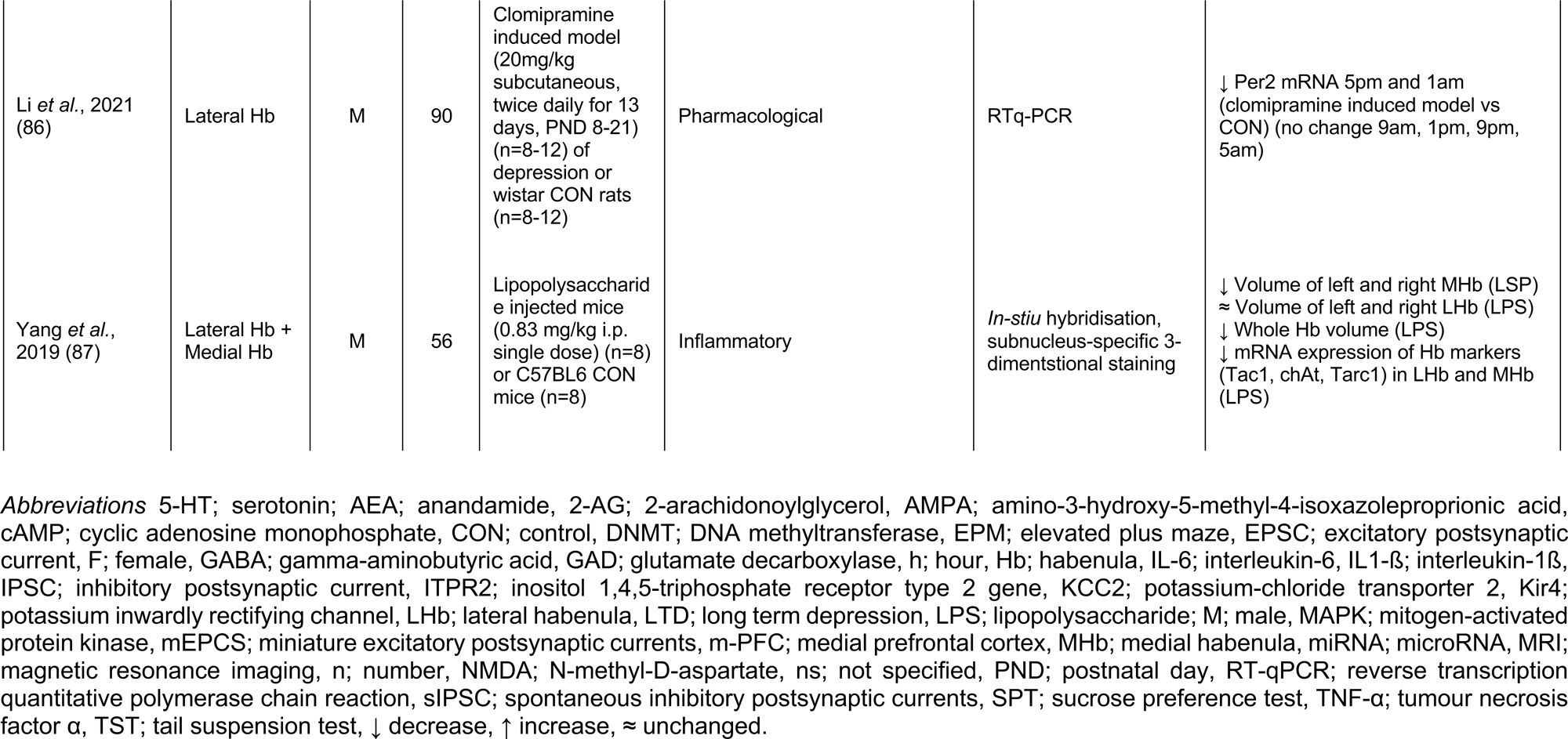
Molecular measures in the habenula across preclinical models of depression.

### 2.1 Search Strategy

A literature search was completed using the following electronic databases; PubMed, Scopus, and Web of Science. All publication dates until Aug 2023 were included in the literature search. The two search terms depress* and habenula* were applied to each of the relevant databases in the field of title, abstract, and/or keyword. For example, the search strategy used for PubMed was (depress* AND habenula*).

### 2.2 Eligibility Criteria

Studies eligible for inclusion in this systematic review must have been full-text, primary, research articles published in English and examined changes in the Hb in clinical cases of depression (living or post-mortem) or preclinical models relevant to depression. Any measure of the habenula was eligible for review including structural, functional, cellular, molecular, and electrophysiological measures. Preclinical articles were excluded if they examined differences in models of depression following the administration of an external treatment or following external manipulation of the Hb, without including a baseline measure in the model compared to control. Clinical studies were excluded if they did not have a non-psychiatric control group or examined depression amongst other diagnoses, where depression was not an independent factor.

### 2.3 Data Selection

In adherence to the PRIMA guidelines, articles obtained from the initial electronic database search were screened by title and abstract by one author (SC). Duplicates were then removed using the reference management software Zotero (Version 6.0.18, Roy Rosenzweig Centre for History and New Media, Virginia, United States). Relevant articles were assessed by full-text based on the inclusion and exclusion criteria to determine their eligibility for review. Where there was ambiguity surrounding the articles, all authors discussed until consensus was reached. The reference list of eligible studies was also screened to identify additional studies.

### 2.3 Data Extraction and Analysis

The following data was extracted from articles that adhered to the inclusion criteria and were determined eligible for review: author, publication date, sample size, subject demographics, and medication usage (for clinical studies only), age, sex (and for clinical studies analysis of sex as a biological variable), the region of habenula examined (medial/lateral), preclinical model used (for preclinical studies only), methods used to obtain habenula measures, and the key findings of the research. Data extraction was recorded using Microsoft Excel (Version 16.68, Microsoft, Washington, United States) and Zotero (Version 6.0.18, Roy Rosenzweig Centre for History and New Media, Virginia, United States) was used to collate references. The current review includes both animal and human studies; accordingly, no risk of bias assessment was performed due to the heterogeneity of study designs and the lack of standardised reporting requirements. In addition, due to the lack of empirically validated risk of bias tools in animal studies, methodical quality assessment was not justified (31).

## 3.0 Results

The primary database search yielded a total of 1648 articles from PubMed (485 items), Scopus (646 items) and Web of Science (517 items) (***Figure 2***). One additional record was identified though the screening of reference lists. Following the exclusion of duplicates, a total of 637 articles were screened by title and abstract for eligibility, 526 articles were excluded as they were not primary research articles, or they did not assess alterations in habenula measures in the clinically depressed population or preclinical models relevant to depression. 111 articles passed the initial screening process and were assessed by full-text review for eligibility. A further 24 articles were excluded because they did not examine baseline changes in preclinical models of depression prior to external manipulation of the Hb, three articles were excluded as they did not independently examine depression within the psychiatric cohort (grouped with schizophrenia, bipolar or other disorders) and nine articles were excluded due to the lack of a non-psychiatric control group. A total of 75 articles that adhered to the eligibility criteria were included in the qualitative synthesis (n=57 preclinical studies; n=18 clinical studies).

The majority of the preclinical studies examined changes in markers of habenula activity (n=16), neuronal firing (n=21) and neurotransmission (n=13) (**Table 1**). There were an additional 7 studies that assessed other Hb measures related to volume and connectivity (n=1), inflammation (n=2), genomic changes (n=2) and circadian rhythm (n=2). Of the 57 preclinical studies, 7 experimental designs (12%) included both male and female animals. From these, 5 studies (71%) reported a significant difference between the sexes in at least one measure taken. Most preclinical investigations were conducted on adult animals (n=43), with some inclusion of infants (n=2), juveniles (n=5) and late adolescents (n=7). In addition, the majority of preclinical studies specifically analysed the LHb, with only 13 preclinical studies measuring changes in the MHb (**Table 1**).

Clinical studies included for review (n=15 living; n=3 post-mortem) measured changes in Hb functional connectivity (n=11), volumetric differences (n=5) and molecular alterations (n=2) (**Table 2**). Of these 18 clinical studies, 16 examined the whole Hb complex and did not differentiate the medial and lateral subdivisions. Clinical studies generally included male and female subjects (n=15), however, few of these studies examined sex as a biological variable (n=5). Instead, sex was typically examined as a covariate and male and female data were pooled together (n=10). In addition, all clinical studies were conducted in adults with the exception of 1 study that examined adolescents.

**Table 2.**
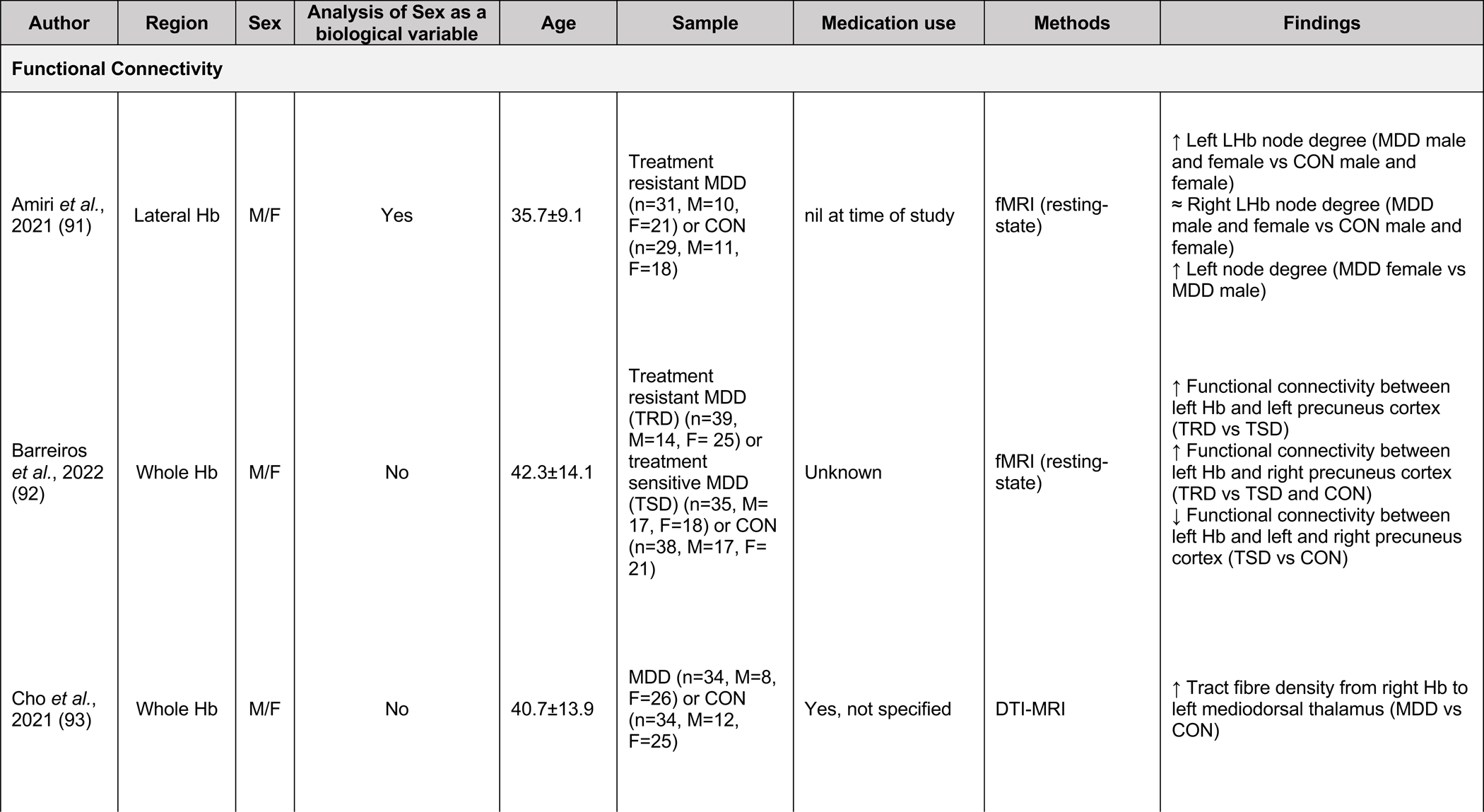
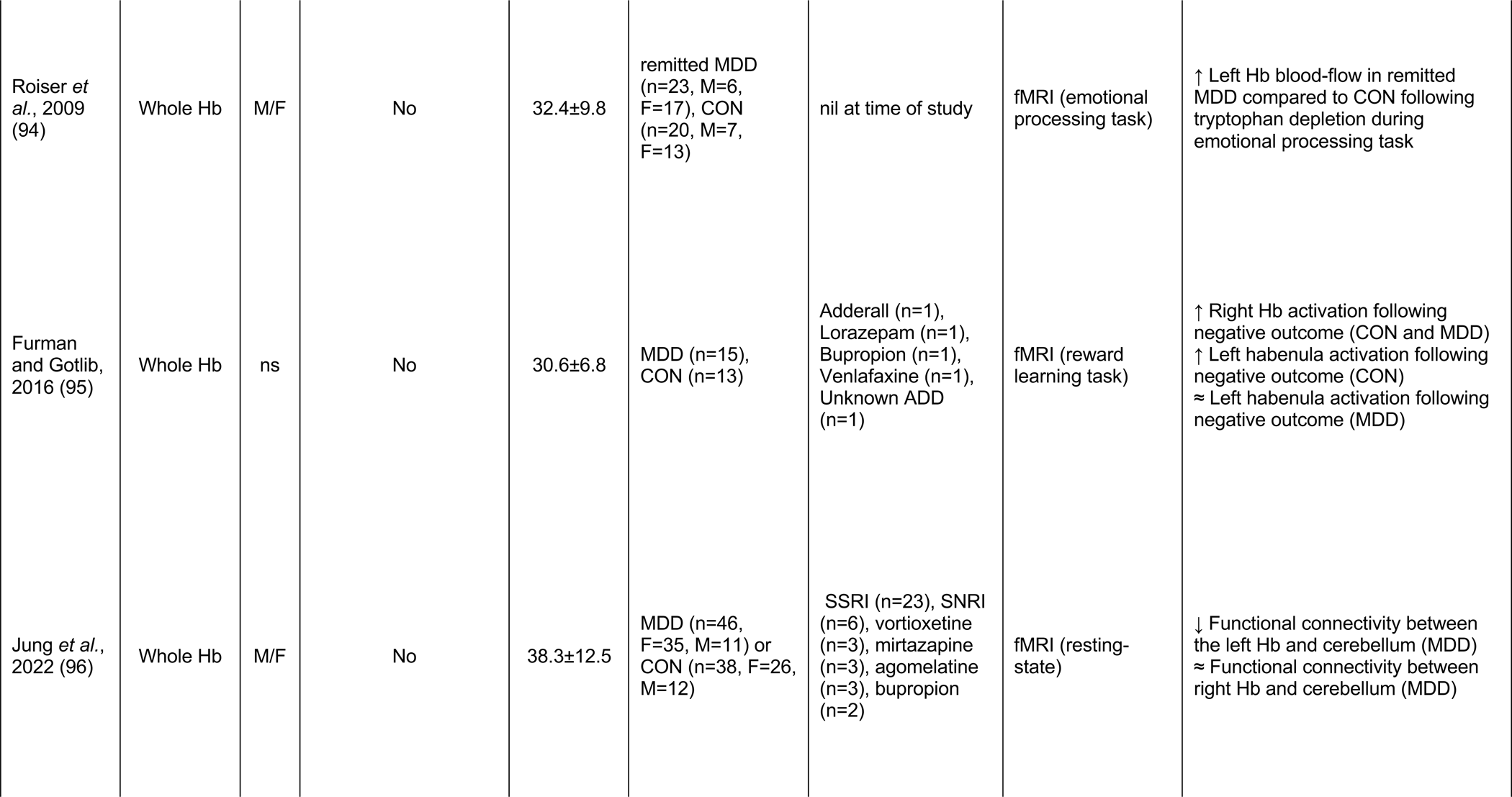
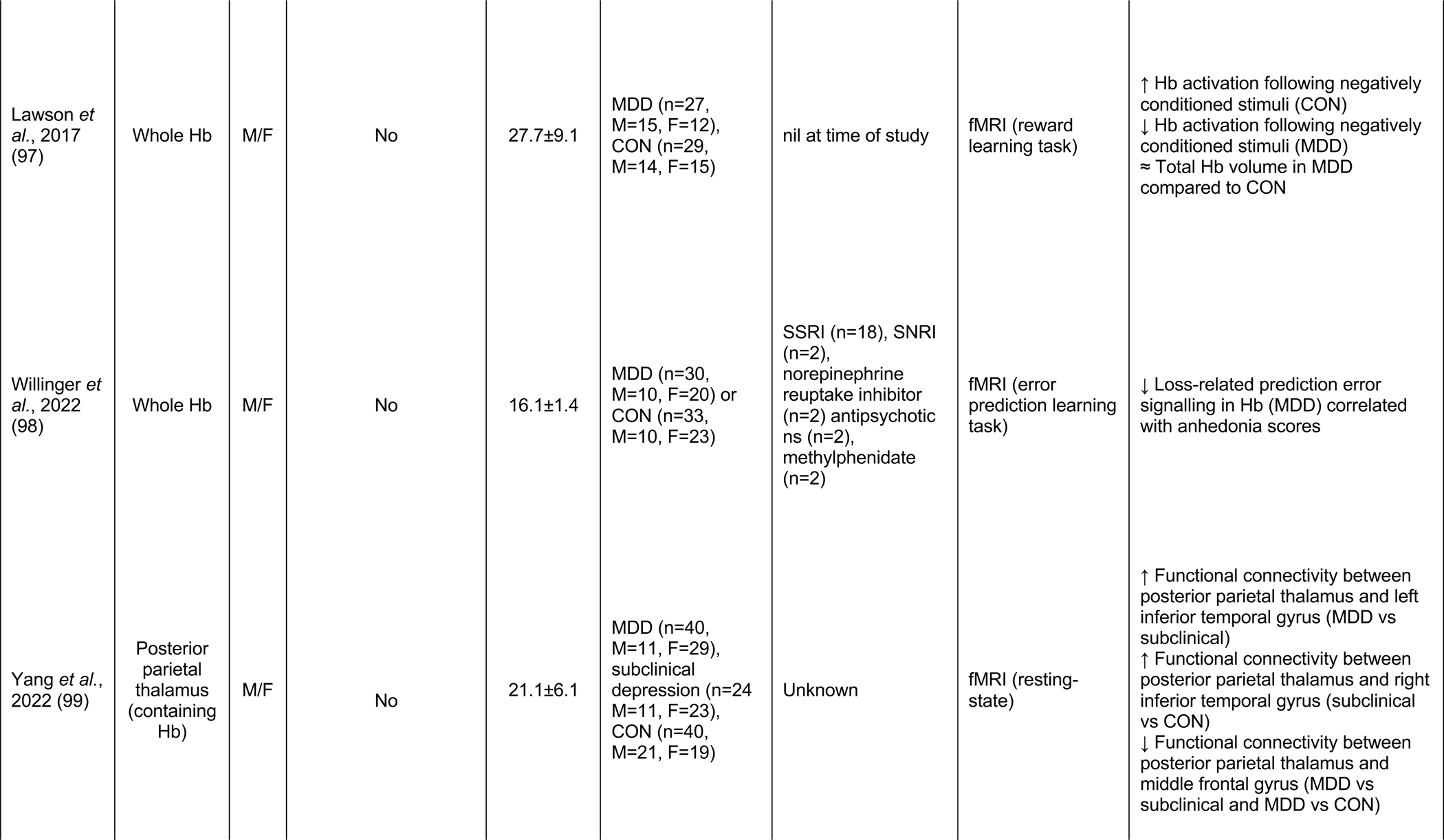
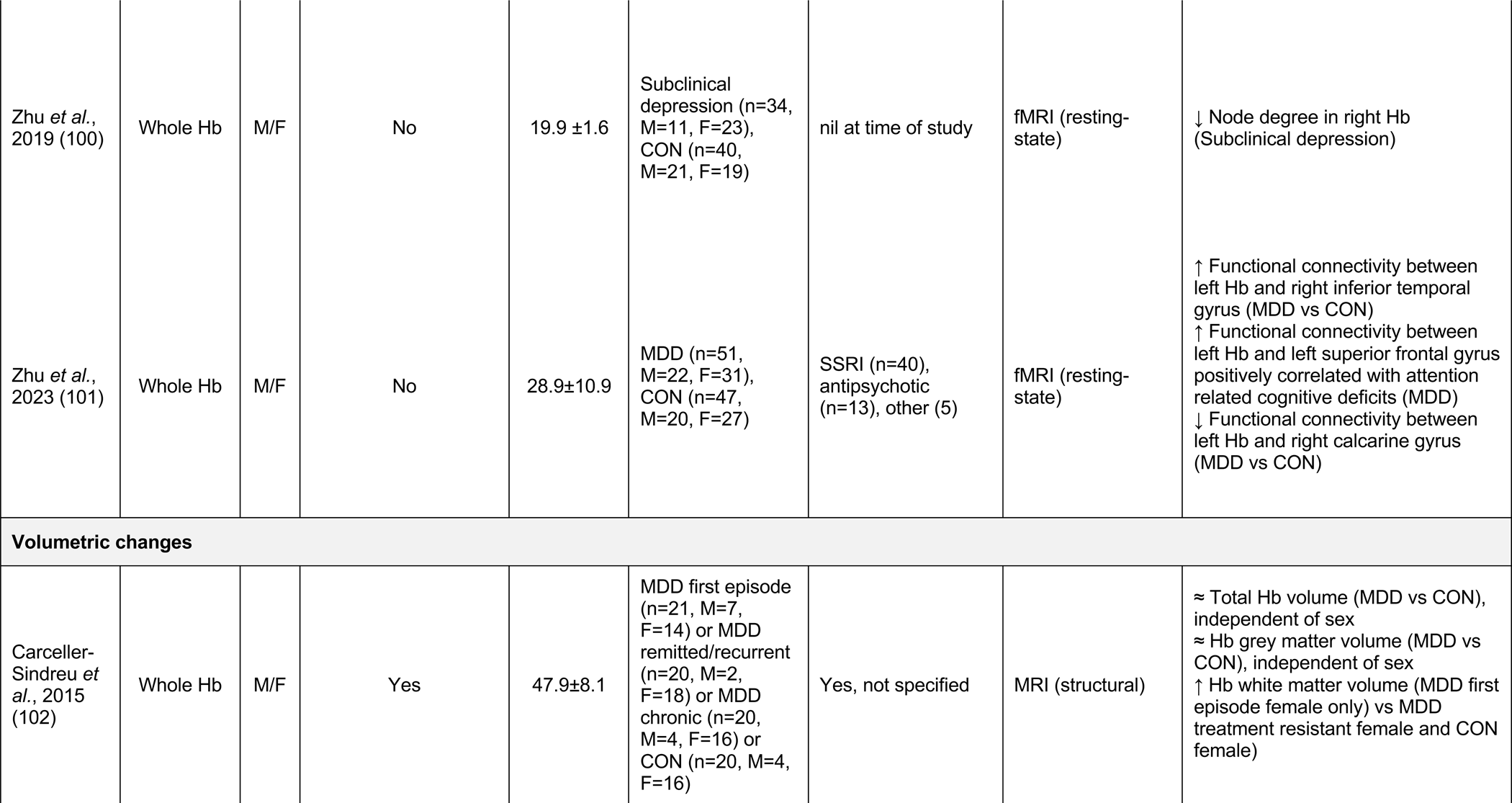
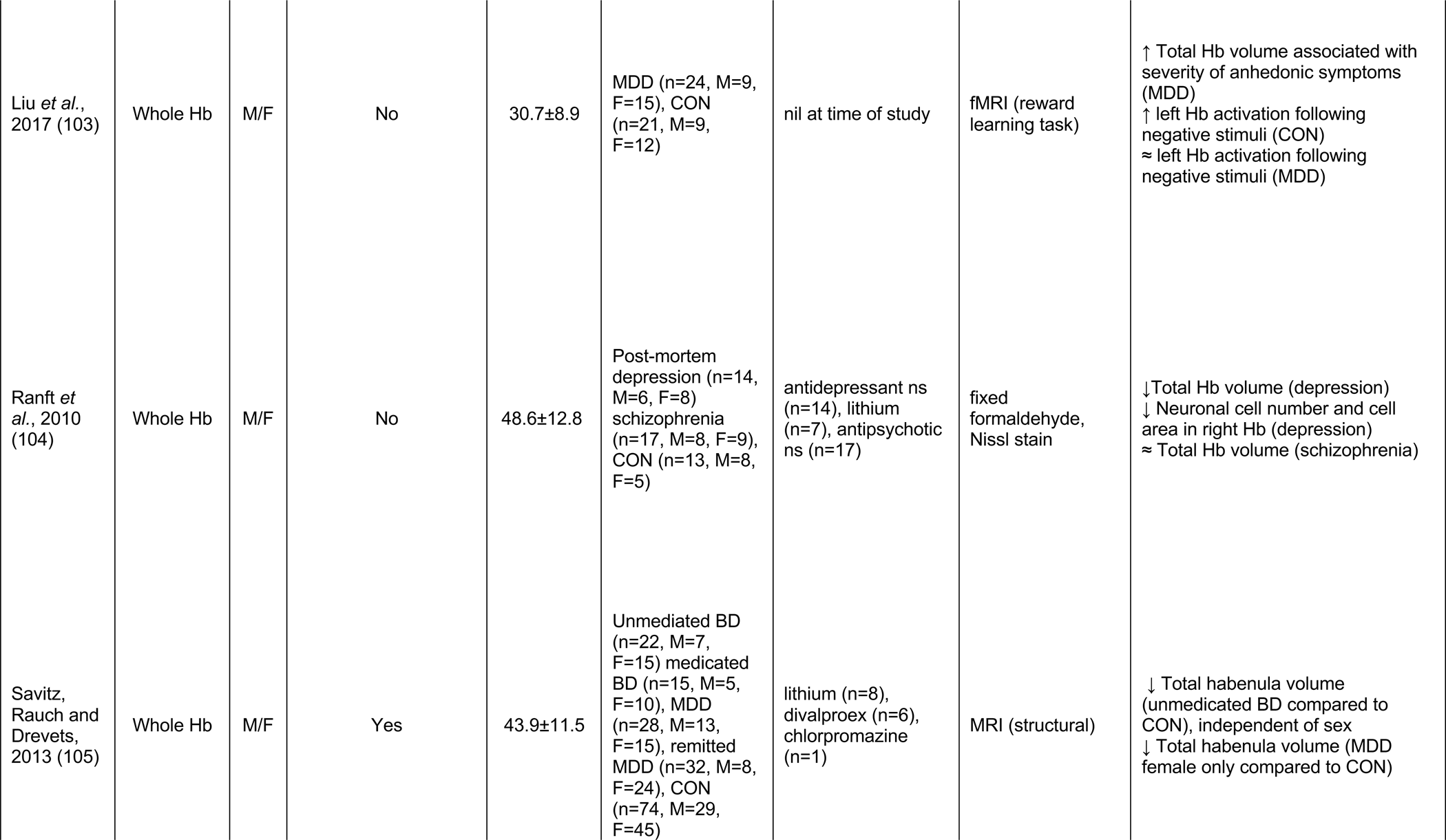
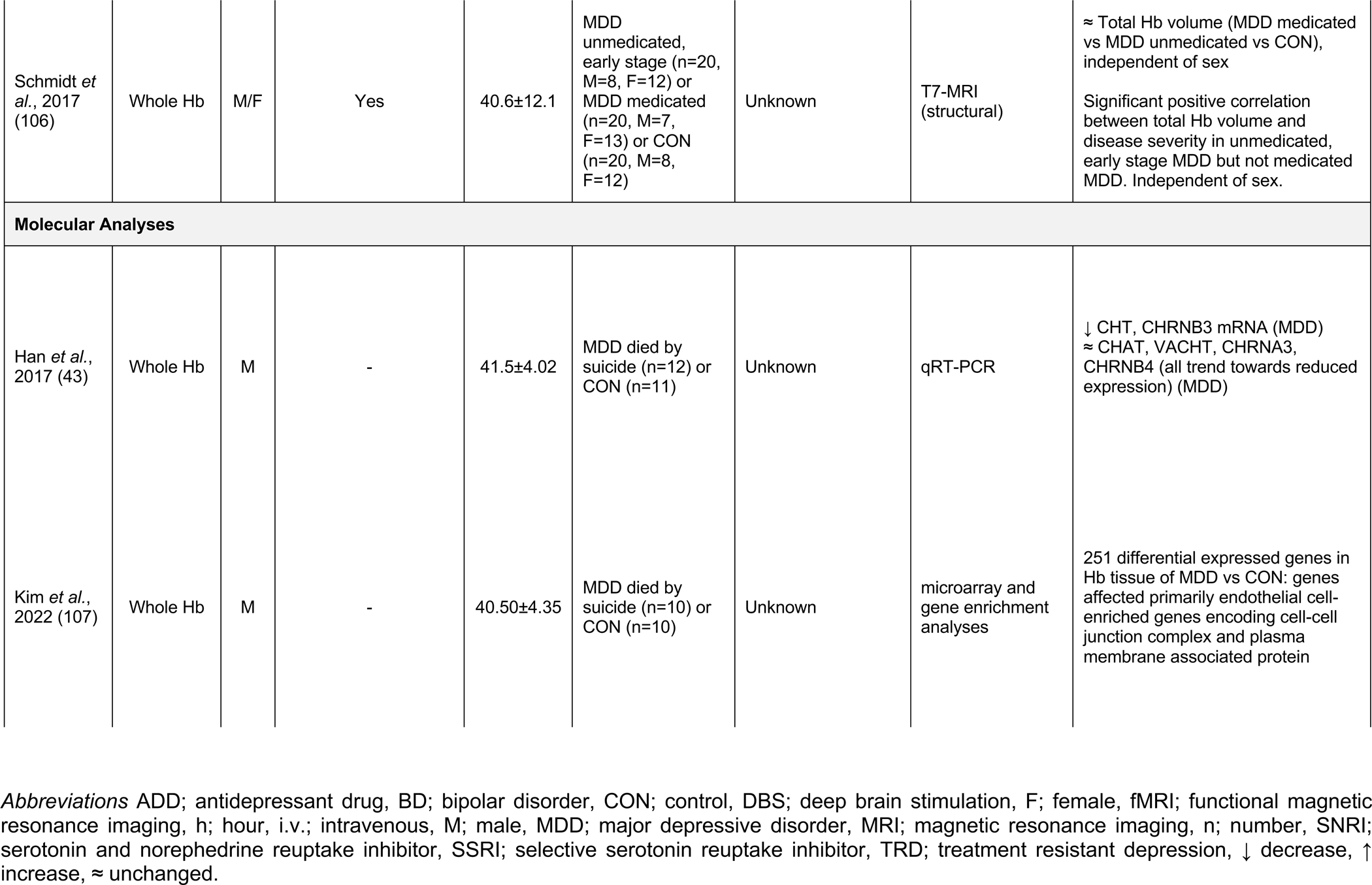
Habenula measures in clinical cases of depression.

### 3.1 Alterations in the habenula across preclinical models of depression

Depressive-like states in preclinical models are commonly induced via exposure to aversive stress. Despite varying in type, duration, and life stage the stressor occurs, these environmental models typically induce behavioural changes analogous to the depressive phenotypes observed in the clinical population. In the current review, the majority of preclinical studies employed an environmental stress paradigm (n=45). Within this group, over 37 different stress protocols were used. Genetic models of depression largely relied on the selective breeding of stress susceptible rodents (n=7) or insertion/knockout of depressive-related genes (n=1) to induce behavioural changes. Chemical-induced models of depression depended on the use of an external treatment and generally provided a more mechanistic approach to understanding the disorder. The initiation of depressive-like states via liposaccharide treatment (inflammatory agent) was the most common chemical-induced model included for review (n=3). Across the preclinical models, evidence suggests Hb hyperactivity, caused by a combination of altered glutamatergic and GABAergic signalling, astrocyte dysfunction and neuroinflammation (***Figure 3***).

**Figure 3.**
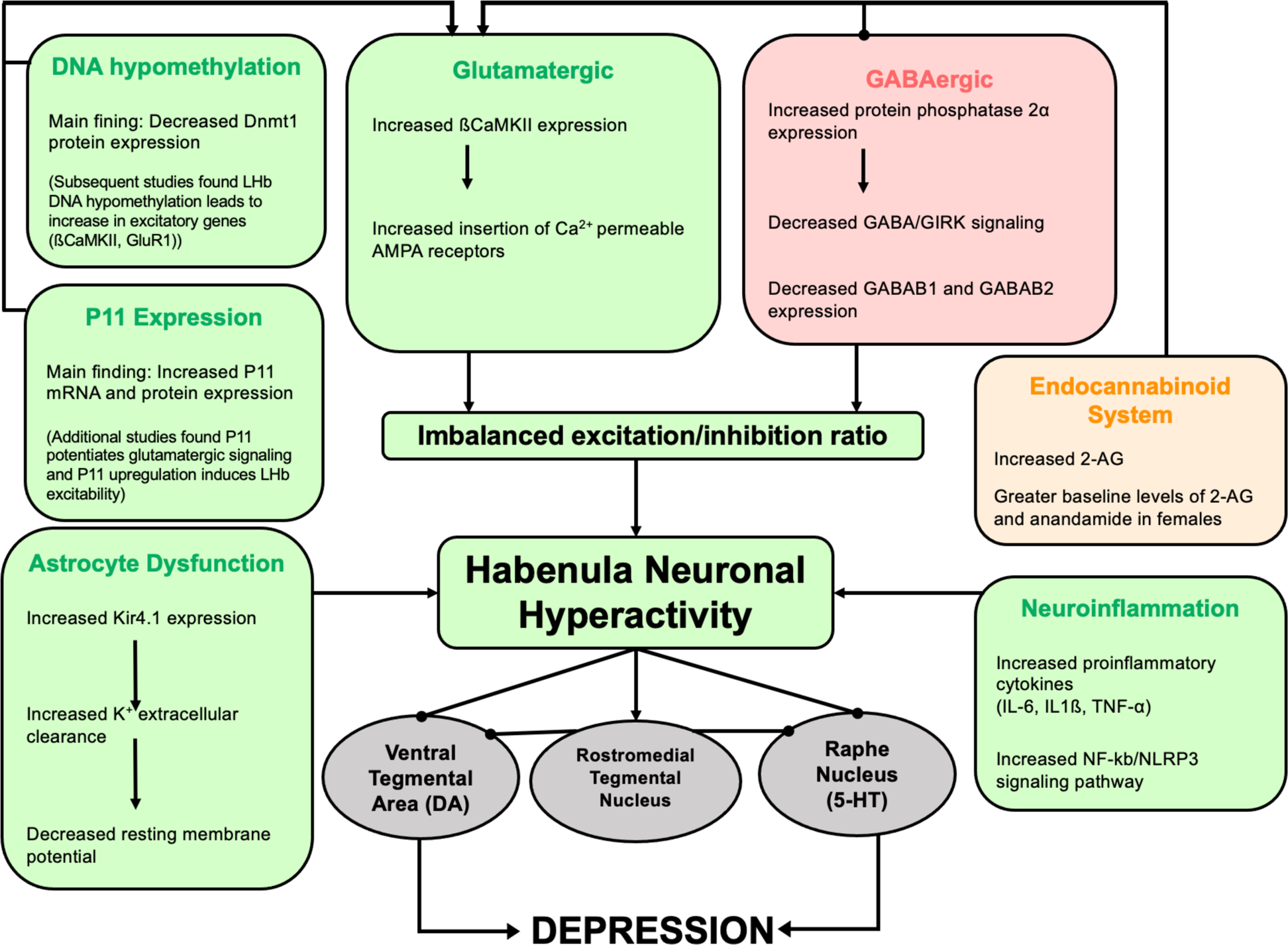
Flow chart summarising the molecular changes that have been reported in the Hb in preclinical models of depression. Excitatory changes are shown in green, inhibitory changes are shown in red and the endocannabinoid system, a neural system influencing both inhibitory and excitatory systems, is shown in orange. Boxes in grey illustrate the efferent projections of the Hb ***Abbreviations*;** AEA, anandamide; AMPA, α-amino-3-hydroxy-5-methyl-4-isoxazoldpropionic acid; ßCaMKII, ß calmodulin-dependent kinase II; DA, dopamine; DNA, deoxyribonucleic acid; Glu, glutamate; GABA, gamma aminobutyric acid; IL-6, interleukin-6; IL-1ß, interleukin-1ß Kir4.1, inward rectifying potassium 4.1 channel; TNF-α, tumour necrosis factor α; 2-AG, 2-arachidonoylglycerol; 5-HT, serotonin; ® activation; inhibition.

#### 3.1.1 Markers of habenula neuronal and metabolic activity

Across preclinical models of depression, markers of neuronal and metabolic activity were consistently upregulated in the habenula (n=16). The neuronal activation marker, c-Fos, was increased in the LHb following exposure to acute restraint stress (63), chronic restraint stress (47, 48), chronic inescapable shock stress (32), chronic unpredictable stress (41, 46) and social defeat stress (53, 56, 57). Only two of these studies examined c-Fos expression in male and female rodents; both studies observed sex differences in the Hb (48, 63). While both male and females displayed an elevation in c-Fos expression following exposure to chronic restraint stress, the increase was significantly greater in the LHb of stressed females compared to stressed males (48). Conversely, following acute stress exposure, Sodd et al. (63) reported a significantly higher c-Fos expression in stressed males compared to females, suggesting sex differences warrants further investigation.

A total of 4 studies reported that Hb neuronal activity did not significantly differ between control and depressive-like rodents; however, each of these studies supported a general trend towards greater Hb activity within the stressed cohort (39, 57, 64, 65). One study found that c-Fos cell-count was unchanged in the MHb and LHb of rats exposed to chronic restraint stress (39). Despite this, an increase in c-Fos staining intensity was observed in animals susceptible to stress compared to those deemed resilient (39). Exposure to social defeat stress in balb/c mice also did not alter c-Fos activity (24h post-stress) (57). However, the stress did lead to a delayed increase in c-Fos activity in the LHb 2 weeks following the initial stress exposure; a similar trend was observed within the MHb (57). Similarly, c-Fos was not changed in the MHb following exposure to social isolation stress; however, a positive trend was noted. (65).

The enzyme, cytochrome oxidase, is an indicator of brain metabolic capacity and greater expression has been shown to correspond to a greater degree of neuronal activity (88). In congenitally learned helpless rats, an increase in cytochrome oxidase was noted in both the MHb and LHb when compared to non-helpless controls (83). Greater Hb cytochrome oxidase expression was also reported in a model of chronic unpredictable stress (71, 33). However, a study conducted by Spivey et al. reported that early life stress, involving either exposure to early handling or maternal separation, did not alter the cytochrome oxidase activity in the LHb (64), suggesting developmental stress may have less effect on Hb metabolic activity. Despite these findings, female rats recorded greater baseline cytochrome oxidase activity in the LHb compared to males, irrespective of stress or control group, further supporting sex differences in Hb activity (64).

#### 3.1.2 Habenula neuronal firing

The most consistent finding across preclinical models of depression was aberrant neuronal firing of the Hb, specifically hyperactivity within the LHb. LHb hyperactivity was reported across 11 experimental designs that utilised 14 models of depression. Rather than abnormal tonic firing, the dysregulated Hb neuronal activity appears to be specific to a subtype of burst neurons. Greater spontaneous bursting activity in the LHb has been reported following chronic (n=7), acute (n=1) and early life stress paradigms (n=3) (42, 50, 52, 62, 51, 61, 75, 84). Similarly, this phenomenon has been observed in the genetic congenitally learned helplessness model (n=2) (72, 79) and in an inflammatory model of depression (42). Moreover, general baseline Hb neuronal activity was greater in the Hb of Wistar Kyoto rats (endogenous treatment-resistant model) when compared to the non-depressive-like hypertensive rat (85).

Exposure to stress is theorised to lead to increased sensitivity within the cells of the LHb, thereby leading to more frequent firing. This is supported by work showing that a lower rheobase (minimum frequency required for an action potential) was recorded in excitatory neurons projecting from the m-PFC to the LHb in both a chronic stress paradigm and a genetic knockout (*itpr*2 knockout) model (54). LHb outputs, specifically to the dopaminergic VTA, have a similar increase in excitatory postsynaptic currents (52) and overall firing rate (77) in models of acute learned helplessness, congenitally learned helplessness (52) and chronic unpredictable mild stress (77). More recently however Cerniauskas *et al.* 2019 showed that chronic stress-induced LHb-VTA bursting activity was specific to animals that showed greater immobility in the tail suspension test (TST), but not those that showed anhedonic behaviour in the absence of immobility, suggesting LHb hyperactivity may be specific to a learned helplessness or passive coping phenotype (36). Similarly, early life stress via maternal deprivation, which induced only a mild depressive phenotype with a modest reduction in sucrose preference and no change in mobility (OFT), grooming behaviour (splash test) or time spent in open borders (OFT), was associated with LHb hypoactivation, again suggesting LHb hyperactivity may be specific or more pronounced in specific behavioural adaptations (70).

#### 3.1.3 Excitatory/inhibitory neurotransmission balance

Alterations in Hb neurotransmission have been briefly characterised in models of relevance to depression, largely focusing on aspects of excitatory and inhibitory neurotransmission. To date, studies have largely focussed on glutamatergic dysfunction (n=6), the primary neurotransmitter of the LHb. Stress-induced models consistently potentiate LHb glutamatergic transmission, shifting the excitatory/inhibitory (E:I) balance towards excitation (36, 50 69). Both stress-induced and genetic models of depression have reported significant elevations in ßCaMKII (which enhance glutamate activity) in the LHb (n=2) (76, 81). In line with the role of ßCaMKII to enhance glutamatergic synaptic transmission via AMPAR phosphorsphoylation and insertion into the synapse, the post-synaptic expression of calcium permeable AMPARs was increased following chronic stress exposure (36).

Dysfunction within upstream regulators of glutamatergic transmission have also been reported in preclinical models of depression (n=2) (59, 60) P11 is a multifunctional protein that acts to promote glutamatergic (and serotonergic) signalling (66). Seo *et al.* 2018 (59) reported an increase in p11+ cells, p11 protein and mRNA expression within the LHb following exposure to chronic stress. The increase in p11 was accompanied by a long-lasting increase in neuronal excitability within the LHb (59). Furthermore, DNA hypomethylation, via a reduction in the DNA methylation marker, DNMT1, has been reported in the LHb following chronic stress (60). Interestingly, the authors reported that inducing local LHb DNA hypomethylation led to an increase in glutamatergic excitatory genes (ßCaMKII, GluR1) suggesting DNA hypomethylation may be a possible upstream mechanism contributing to hyperactivity within the region (60).

Disrupted inhibitory signalling has been briefly reported in environmental stress models (n=2). With mice susceptible to stress following chronic social defeat showing lower GABA B1 and B2 expression in the LHb (53). In contrast, the MHb displayed a reduction in GABA B1 and B2 following chronic social defeat irrespective of whether the mice were resilient or susceptible to that stress, further highlighting the functional differences between the medial and lateral subdivisions (53). In addition, the inhibitory somatostatin 2 receptor, was increased in the MHb following chronic mild stress exposure (the LHb was not examined) (38).

Dysfunctional endocannabinoid signalling has been reported in one study design. The endocannabinoid system is a neuromodulatory system that regulates glutamatergic and GABAergic signalling at both the pre- and post-synaptic terminals (34). Exposure to acute, chronic, and social defeat stress resulted in an increase in the endocannabinoid, 2-AG, that was correlated to greater LHb neuronal activity, an effect that was independent of sex (34). While stress exposure had no significant effect on anandamide levels, control females recorded higher baseline anandamide concentrations compared to males (34).

#### 3.1.4 Other molecular changes in the habenula

The inflammatory hypothesis of depression has gained momentum in recent years and inducing inflammation is an established preclinical model of depression (87). Interestingly, following systematic administration of an inflammatory agent, Yang *et al.* (87) reported that depressive-like behaviours were accompanied by a significant reduction in volume of the whole Hb complex. When accounting for the different divisions it was determined this change was predominantly driven by changes within the MHb not LHb (87). Inflammatory changes are also described as a secondary response to stress exposure and two studies have reported an elevation in inflammatory cytokines within the LHb (44, 69). An increase in TNF-α, IL-1ß and IL-6 was noted in a model of chronic unpredictable stress and was accompanied by greater activation within the inflammatory NF-kB/NLRP3 signalling pathway (69). In addition, following exposure to chronic social defeat stress, mice were separated into a susceptible group that exhibited depressive-like behaviours and a resilient group that did not (44). Compared to non-stressed control mice, both susceptible and resilient mice displayed an increase in the pro-inflammatory markers TNF-a, IL-1ß and IL-6 in the LHb (44). Stress exposed mice also displayed greater activation within the matrix metalloproteinase (MMP) 12-MMP2 pathway, however; an increase in Pcsk5 (MMP upstream regulator that facilitates microglia motility) and greater monocyte mobilisation was specific to the stress susceptible group (44). Ito et al. suggest that greater cytokine release may lead to a remodelling of the extracellular matrix providing a possible explanation for these findings (44).

A comprehensive genomic profile of the Hb in rats exposed to chronic stress revealed the differential expression of 379 genes (73). Of these, stress exposure particularly affected cholinergic synapse-related genes associated with neuronal signalling in the MHb (73). In addition, downregulation of several serotonergic receptor genes was reported in stressed rats compared to non-stressed controls (73). Serotonergic dysfunction has also been reported in the Hb of the Wistar Kyoto rat, an endogenous model of treatment resistant depression (89), with downregulation of the serotonergic receptor genes *Htr7* and *Htr2a* in both the MHb and LHb and increased expression of *Htr4* in the LHb but not MHb (80). Similarly, a reduction in 5-HTR7 receptor expression was isolated to the LHb (80).

Disruptions in circadian rhythm have previously been linked to depression in both clinical and preclinical investigations (reviewed in 90). The Hb is situated near the pineal gland, a prominent regulator of the circadian cycle. Consequently, Hb dysfunction has also been examined in the context of sleep, specifically clock genes that contribute to the homeostatic aspect of sleep regulation (n=2) (37, 86). Christiansen et al. (37) reported a reduction in the *Per2* clock gene in the LHb of rats exposed to chronic stress. The down-regulation of *Per2* has previously been noted in other brain regions that are hypoactive in depression and is suggested to contribute to depressive symptoms at night (37). Similarly, in a clomipramine induced (20 mg/kg subcutaneous) model of relevance to depression, *Per2* mRNA was significantly reduced in the LHb compared to controls, in a time-dependent manner (86).

### 3.2 Alteration in habenula measures in clinical depression

Clinical research is limited by the accessibility of the Hb, leading studies to predominately rely on non-invasive neuroimaging methods. The majority of the literature examined basal changes in Hb connectivity in cases of clinical depression (n=8) or during task-based conditions (n=3). Volumetric and morphological changes in the Hb were also assessed (n=5). A further two studies conducted post-mortem molecular analyses of the LHb. Subtypes of depression examined were limited to major depressive disorder (n=14), treatment resistant depression (n=2) and subclinical depression (n=1). In addition, as expected, subjects with depression in nearly all included studies had some history of anti-depressant drug use (n=16).

#### 3.2.1 Resting-state fMRI

Functional connectivity is correlated to greater cerebral blood flow and is suggested to be indicative of increased metabolic and subsequent neuronal activity (108). A study conducted by Amirir et al. (91) reported that greater Hb node degree (indicative of a higher number of connections) was characteristic of treatment-resistant depression. The greater node degree was specific to the left hemisphere and more pronounced in females compared to males within the same diagnosis group (91). Conversely, in a subclinical depression cohort, left Hb node degree was not different compared to non-psychiatric controls, suggesting the severity of depression influences Hb connectivity (100). A study by Yang *et al.* (99) further supports the correlations between greater functional connectivity of the Hb and disease severity. Compared to controls and those with subclinical depression, individuals with a diagnosis of major depression showed greater functional connectivity between the posterior parietal thalamus (region containing Hb) and the inferior temporal gyrus. Individuals with depression also showed greater functional connectivity between the left Hb and the inferior temporal and superior frontal gyrus but reduced connectivity to the calcarine sulcus (100) and cerebellum (96); no change was recorded in the right Hb. Interestingly, Barreiros et al. (92) identified that patients with treatment-resistant depression exhibited greater functional connectivity between the left Hb and precuneus cortex. Conversely, individuals deemed responsive to traditional anti-depressants showed hypoconnectivity between the left Hb and precuneus cortex when compared to controls (92).

#### 3.2.2 Reward learning task fMRI

Three studies have reported abnormalities in habenula activity in depressed individuals during reward-task based fMRI. Considering the Hb signals negative stimuli, it is expected that Hb activity would increase in response to a negative outcome. Indeed, both left and right Hb activation has been shown to increase in healthy controls when exposed to a negative outcome (95, 96). However, those with a diagnosis of MDD recorded either no or attenuated change in Hb activation following exposure to the same negative stimuli (95, 96). Similarly, in an error prediction learning task, adolescents with MDD exhibited a reduction in negative related signalling in the Hb when compared to controls (98). Moreover, the reduction in loss-related prediction error correlated with anhedonia symptom severity 98). While the reduction in Hb activation appears to contradict the preclinical research, it does support a perturbed habenula response in MDD.

#### 3.2.3 Volumetric changes

Habenula volumetric studies (n=5) have produced inconsistent results, and comparison is difficult due to the various methods used, which include a combination of ante- and post-mortem analyses (102, 103, 104, 105, 106). Ranft et al. reported a total decrease in Hb volume in post-mortem depression cases (n=6) (104). This was accompanied by a reduction in cell number and cell area in the right Hb; a similar though insignificant trend was noted in the left Hb (104). A high-resolution MRI study with a larger sample (n=28; M=13, F=15) reported a similar reduction in total Hb volume, however; these findings only pertained to females with a diagnosis of MDD (105). Interestingly, while Carceller-Sindreu et al. reported no overall difference in whole Hb volume, the study found an increase in Hb white matter volume in female first episode MDD patients compared to female treatment-resistant MDD patients (102). These results did not extend to the male MDD cohort or the Hb grey matter (102). Taken together, these studies highlight the importance of considering sex in studies of the Hb in depression. While two further studies have shown no overall difference in Hb volume in depression compared to control subjects, both report a relationship with symptoms, specifically that an increase in Hb volume was associated with greater depressive symptom severity (103, 106).

#### 3.2.4 Post-mortem molecular changes

One post-mortem study comprehensively investigated differences in gene expression profiles between male MDD subjects that died by suicide and non-psychiatric controls (107). The study found a total of 251 differentially expressed genes in the Hb of MDD patients compared to controls. These genes primarily affected endothelial cell-enriched genes encoding cell-cell junction complexes and plasma membrane associated proteins and their upstream regulators (107). Endothelial dysfunction may compromise blood brain barrier permeability (107). In an additional study conducted by the same research group, individuals with MDD that died by suicide showed a reduction in cholinergic genes (CHT, CHRNB3) in the Hb post-mortem (43).

## 4.0 Discussion

Overall, the developing body of evidence consistently points to aberrant Hb neuronal firing and dysregulated Hb neurotransmission, particularly excitatory/inhibitory balance, in preclinical models of depression. The hyperactivity observed within preclinical models translates to greater functional connectivity observed within a subgroup of Hb projections in the clinical population (91, 93, 95). The small number of clinical studies that have examined molecular changes in the Hb, shed light on previously unexplored mechanisms in preclinical models of depression including endothelial dysfunction and compromised blood brain barrier integrity. A complete summary of the proposed molecular mechanisms underlying Hb dysfunction in preclinical models of depression and the clinical population is illustrated in ***Figure 4***.

**Figure 4.**
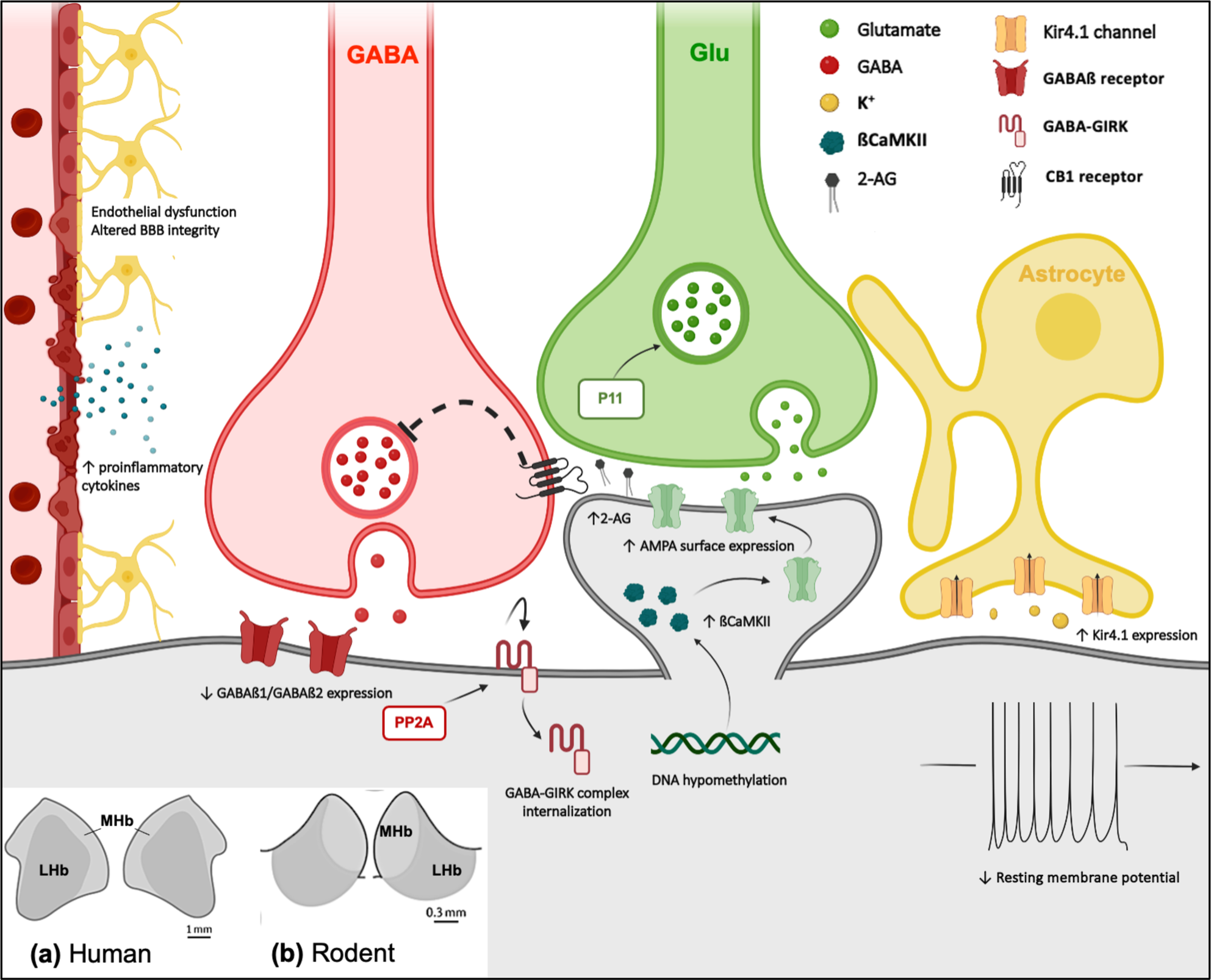
Schematic representation of the potential molecular mechanisms contributing to Hb hyperactivity in preclinical models of depression and the clinical population. DNA hypomethylation leads to an increased expression of glutamatergic excitatory genes. Greater ßCaMKII gene and subsequent protein expression then results in an increased surface expression of AMPAR. Together with increased p11 protein expression, these molecular changes potentiate glutamatergic signalling. GABAergic transmission is attenuated via reduced expression of GABAß1/2 and increased activity of the PP2A enzyme which leads to GABA-GIRK complex internalisation. Increased expression of the endocannabinoid, 2-AG, acts via the CB1 receptor to inhibit GABA release. Greater expression of the Astrocytic Kir4.1 potassium transporter, reduces the resting membrane potential leading the neuron in a state more primed to fire and compromised BBB integrity leads to greater cytokine infiltration. A comparison of the **(a)** human and **(b)** rodent Hb and the respective boundaries of the MHb and LHb is shown in the bottom left. ***Abbreviations***; AMPA, α-amino-3-hydroxy-5-methyl-4-isoxazoldpropionic acid; ßCaMKII, ß calmodulin-dependent kinase II; BBB, blood brain barrier; CB1, cannabinoid type 1 receptor; Glu, glutamate; GABA, gamma aminobutyric acid; Kir4.1, inward rectifying potassium 4.1 channel; LHb, lateral habenula; MHb, medial habenula; PP2A, protein phosphatase 2A; VHb, ventral habenula; 2-AG, 2-arachidonoylglycerol

### 4.1. LHb: an emerging model of altered excitatory/inhibitory balance in preclinical models of depression

A clear consensus emerges from the preclinical research linking aberrant Hb neuronal firing to the development of depressive-like behaviours. Enhanced glutamatergic transmission combined with attenuated inhibitory signalling may lead to a state of abnormal activation, specifically within the LHb. Findings from the MHb generally support this pattern, however, considering the diversity in neuronal populations between the two subregions further research is necessary. Subsequent advancements in optogenetics have allowed researchers to determine whether Hb hyperactivity plays a causative role in the development of depressive-like behaviours. Indeed, in healthy rodents, direct stimulation of the LHb is sufficient to induce a depressive-like phenotype, suggesting LHb hyperactivity may be a primary pathology in depression (22, 49). Preclinical evidence suggests molecular substrates involved in glutamatergic transmission in the Hb may be a valuable target to suppress neuronal firing (50, 45, 76, 81). Overexpression of the enzyme, ßCaMKII, which promotes glutamatergic transmission, in the LHb induces symptoms of behavioural despair and anhedonia whereas blockade of ßCaMKII ameliorates these depressive-like behaviours (81). Moreover, the antidepressants impramine and ketamine downregulate ßCaMKII levels in the LHb (76, 81).

Enhanced glutamatergic transmission may be the primary driver of Hb hyperactivity; however, dampened GABAergic signalling seems to further offset the Hb excitatory/inhibitory balance (50, 53, 68). Expression of GABA(B) receptors is downregulated in mice susceptible to stress (53). Moreover, GABA(B)-GIRK complex internalisation within the LHb has been implicated in the development of depressive behaviours (51). Interestingly, treatment with both a GABA(B) agonist and antagonist produced anti-depressant like effects (53). The localisation of GABA(B) receptors in the Hb is unclear, and it may be that they are located on both the pre- and post-synaptic terminals (53). An important consideration of these treatments is first identifying the location of each receptor and whether the drugs differ in their affinity for the differentially located subtypes (53). The endocannabinoid system is an additional neural system responsible for regulating glutamatergic and GABAergic transmission (34). Results reported by Berger et. al. suggest an increase in endocannabinoids, a mechanism intended to aid in neuronal homeostasis, further accentuate neural imbalances in the Hb (34). Rather than suppressing neuronal excitability, as previously noted in cortical and hippocampal brain regions (110), an increase in the endocannabinoid, 2-AG, further facilitates neuronal activation in the Hb (34). The paradoxical results may be explained via the endocannabinoids preferential suppression of GABA, via its potentiation of glutamate or via differing actions on the pre- and post-synaptic terminals (109). Understanding how this applies in the human brain, including the localisation of cannabinoid receptors on glutamatergic or GABAergic terminals in the Hb, will be an important step forward.

An additional driving regulator of LHb bursting activity involves glial-neuron interactions. The potassium channel, Kir4.1, expressed on astrocytes regulates the degree of membrane polarisation (78). Overexpression of astrocytic Kir4.1 in the LHb induces neuronal bursting that is reversed via ablation of Kir4.1 (78). Moreover, preclinical models of depression show greater expression of Kir4.1 in the LHb (78). Upregulation of Kir4.1 leads to an increased clearance of extracellular K^+^ (78). The lower levels of extracellular K^+^ then induce membrane hyperpolarisation which may explain the abnormal firing pattern (78) Maladaptive astrocyte functioning and subsequent disrupted glutamate clearance has been associated with depressive states and may contribute to abnormal Hb activity (reviewed in 111). Moreover, astrocytes play a vital role in neuroimmune response and are capable of both mitigating and promoting inflammatory signalling (112). While astrocytes are known to be abundant in the healthy Hb, changes in their expression are yet to be examined in the context of depression (113)

Neuroinflammation is an emerging hypothesis underlying depression. In the LHb, chronic stress exposure has been shown to lead to an increase in pro-inflammatory cytokines (44, 69). Local injection of TNF-α into the LHb was sufficient to induce depressive-like symptoms in rodents (69). Moreover, administration of the anti-inflammatory agents aspirin or a NF-kB inhibitor restored these behavioural deficits (69). Interestingly, these findings did not translate when administered locally within the DRN or paraventricular nucleus, suggesting a specific role for the LHb in the inflammatory response (69). Lesions to the LHb also reduced the expression of proinflammatory cytokines induced by chronic stress and improved the hippocampal anti-inflammatory response via activation of the PI3K/mTOR pathway and reduced the expression of apoptosis-related proteins (69). Given the proximity of the Hb to the third ventricle and previous reports of altered blood brain barrier permeability in depression, the region may be particularly vulnerable to cytokine infiltration and be a unique target in the therapeutic action of anti-inflammatory treatments (107).

### 4.2. Clinical insights

Clinical studies support a perturbed Hb response in individuals with depression, the limited evidence points to a general pattern of hyperconnectivity and volumetric changes in depression. However, it is unclear how Hb functionality may vary across sex, age, depression severity and subtype or how external medications may influence its activity in humans. Additionally, clinical studies were typically limited to neuroimaging approaches, which lack the invasive techniques required for accurate measuring of Hb activation. Only two studies have conducted molecular (genetic) analysis on the Hb in people with depression and both studies were limited to males (43, 107). The intriguing results suggest abnormal Hb activity may be a downstream consequence of endothelial cell dysfunction and altered blood brain barrier permeability (107). However, further research is required to understand how these findings integrate into the current hypothesis around Hb dysfunction in depression and determine if the findings are replicable in the female brain.

Considering the variety of networks projecting from the Hb, it is possible that some circuitry may be hyperactive and others hypoactive in depressive states. For instance, while overall Hb activation was consistently elevated in depression, specific connections to the cerebellum and calcarine sulcus were found to be reduced compared to controls (96, 101). Interestingly, a post-mortem human study identified individuals that died with depression displayed a reduction in total Hb volume and cell count (104). While both male and female subjects were included in experiments, sex was not analysed as a biological variable (104). The pooling of sex is an important limitation considering more recent investigations reported Hb volume reductions only pertained to females with a diagnosis of depression (105). Of important consideration, reductions in Hb volume were limited to depression and did not extend to the post-mortem brain of schizophrenia patients (104). Considering schizophrenia often includes depressive symptoms, an interesting avenue for future research involves investigating whether Hb abnormalities are unique to primary depression or occur when depression is secondary to a differential diagnosis such as schizophrenia, dementia, stroke, or Parkinson’s disease.

### 4.3. Therapeutic potential

People with depression often do not experience an alleviation of symptoms until weeks following the initiation of SSRI treatment (6). This delayed therapeutic response aligns with the time required to restore Hb activity in preclinical models (49, 65). In the clinical population, following the administration of single-dose ketamine, subjects with treatment resistant depression exhibited a significant reduction in LHb glucose metabolism compared to baseline, suggesting an attenuating effect on LHb activity (114). While ketamine does not strictly target the LHb, these studies highlight ketamine’s ability to normalise Hb hyperactivity and suggest direct targeting of the Hb may be a promising avenue for future research.

Clinical studies have also suggested alterations within the Hb complex may be indicative of treatment response. In treatment-resistant psychiatric patients (diagnosis of depression, anorexia nervosa or bipolar disorder) receiving subcallosal cingulate deep brain stimulation, individuals unresponsive to treatment had a reduction in total Hb volume at 12 months follow up (115). Conversely, patients that exhibited a therapeutic response displayed an overall increase in Hb volume 12 months post-treatment (115). In addition, high frequency deep brain stimulation of the LHb led to complete remission in a female living with treatment-resistant depression (116) Interestingly, depressive symptoms returned within days following the ceasing of treatment but resolved following the reinstatement of LHb stimulation for 12 weeks (116). It is unclear how deep brain stimulation impacts LHb connectivity and neurotransmission; however, deciphering the underlying mechanisms may help identify the molecular changes in the Hb leading to the antidepressant effect. Furthermore, as previously highlighted, local administration of agents that modulate excitatory (ßCaMKII blockade, NMDA antagonist), inhibitory (GABA B agonist/antagonist) and inflammatory (NF-kB inhibitor, aspirin) processes in the Hb has yielded notable therapeutic outcomes. Examining the molecular composition and cellular architecture of the human Hb may reveal unique treatment targets in this crucial brain region.

### 4.4. Considerations, limitations, and future directions

A major limitation of the current preclinical and clinical literature is the lack of consideration of biological sex. The few studies that conducted molecular analyses on the female Hb demonstrate the results often differ to males. For instance, in rodents while both sexes exhibit LHb hyperactivation, the changes appear to be more pronounced in females following chronic but not acute stress exposure (48, 63). Similarly, females displayed greater basal LHb activation compared to their male counterparts, suggesting they may be particularly vulnerable to changes in Hb firing activity (48, 74). Considering the LHb signals disappointment and aversion, it is theorised that the greater excitation within female mice may bias the animals towards negative emotional states and impair their resilience to stress (117). Whether these molecular sexual dimorphisms contribute to the clinical sex differences observed in affective disorders is yet to be explored. To date, the only molecular characterisation of the human Hb is derived from males and has been conducted at the gene expression level (107). Furthermore, the current literature lacks a fundamental understanding as to whether the cellular, anatomical, and functional composition of the Hb is synonymous between sexes and what the clinical implications of this may be.

Despite emerging evidence highlighting the importance of neurodevelopment on Hb circuitry, the current literature largely focused on the mature, adult Hb. Depression is influenced by both early life environmental factors and genetic predispositions (121). Stressful early life experiences have been shown disrupt LHb maturation, leading to long lasting behavioural changes in adulthood (119). Additionally, specific developmental Hb subtypes has been linked to known risk genes (*Rbfox1* and *Lhx2*) for MDD (120), hinting that changes in the development of the LHb may contribute to the development of MDD. Despite these findings, our understanding surrounding the organisation and development of Hb neurocircuitry remains incomplete (120). It is also unclear when Hb dysregulation occurs during depression onset and how Hb function changes over the course of the disease. At the other end of the spectrum, there is a lack of evidence on Hb functionality in ageing. A critical step forward will be to determine how the properties and function of the Hb alter in the context of the ageing brain, and whether these changes are protective or increase susceptibility to Hb dysregulation and depression in later life.

A limitation that needs to be considered is the small size of the medial and lateral Hb; typical imaging techniques lack the spatial resolution to differentiate the nuclei from one another (122). Consequently, the precise roles and molecular profile of the medial and lateral subdivisions remain elusive in humans. Considering neuroimaging studies are unable to isolate the MHb and the preclinical research has predominately focussed on the LHb, the specific role of the MHb has largely been neglected. This review highlights a need for further consideration of the individualised functions of each subdivision, specifically in the human brain, moving beyond neuroimaging methods and considering post-mortem molecular analyses.

The invasive nature of preclinical research has allowed for comprehensive analysis of Hb neuronal signatures including neurotransmitter populations, neuroconnectivity and cytoarchitecture within rodents (123, 124). However, the neurochemical profile of the human Hb has previously been reported to differ from rodents (21). An essential step in developing the clinical literature is to determine the comparative physiology and cellular organisation (neuronal and non-neuronal cell types, neurotransmitter populations, afferent/efferent projections) of the human Hb. Subsequent investigations should then consider how the Hb may change across human disease states including the various subtypes of depression (bipolar depression, post-stroke depression etc.) and what the implication of these changes may be.

### 4.5. Conclusion

Overall, the preclinical studies presented in this systematic review support Hb hyperactivity as a primary driver for the development of depressive symptoms. The molecular mechanisms driving Hb hyperactivation have been explored and centre on altered inhibitory/excitatory balance, abnormal glial interactions, and neuroinflammation. Moreover, when successful in eliciting a therapeutic response, serval antidepressant treatments including SSRIs, deep brain stimulation, and ketamine intersect in their ability to restore and normalise Hb connectivity, suggesting a role for the Hb as a site for therapeutic agents. However, clinical findings suggest the complexity of depression may go beyond the simplicity of an overactive Hb. Evidence from human studies does support gross Hb abnormalities such as altered activation, connectivity, and volume and as a clinical feature of depression; however, it highlights the need for considerations including hemispheric differences, the specific roles of the medial and lateral subdivisions, and great molecular characterisation of the human Hb in depression. Furthermore, considerations need to be given to the influence of sex and age across preclinical and clinical contexts. Bridging these gaps in the research is essential to enhance our understanding of how Hb dysregulation mediates the core symptoms of depression and shed light on the development of novel therapeutics.

## Declaration of Competing Interest

The authors declare no conflict of interest.

## Acknowledgements

This study has been conducted with the support of the Australian Government Research Training Program Scholarship awarded to S.C.

---

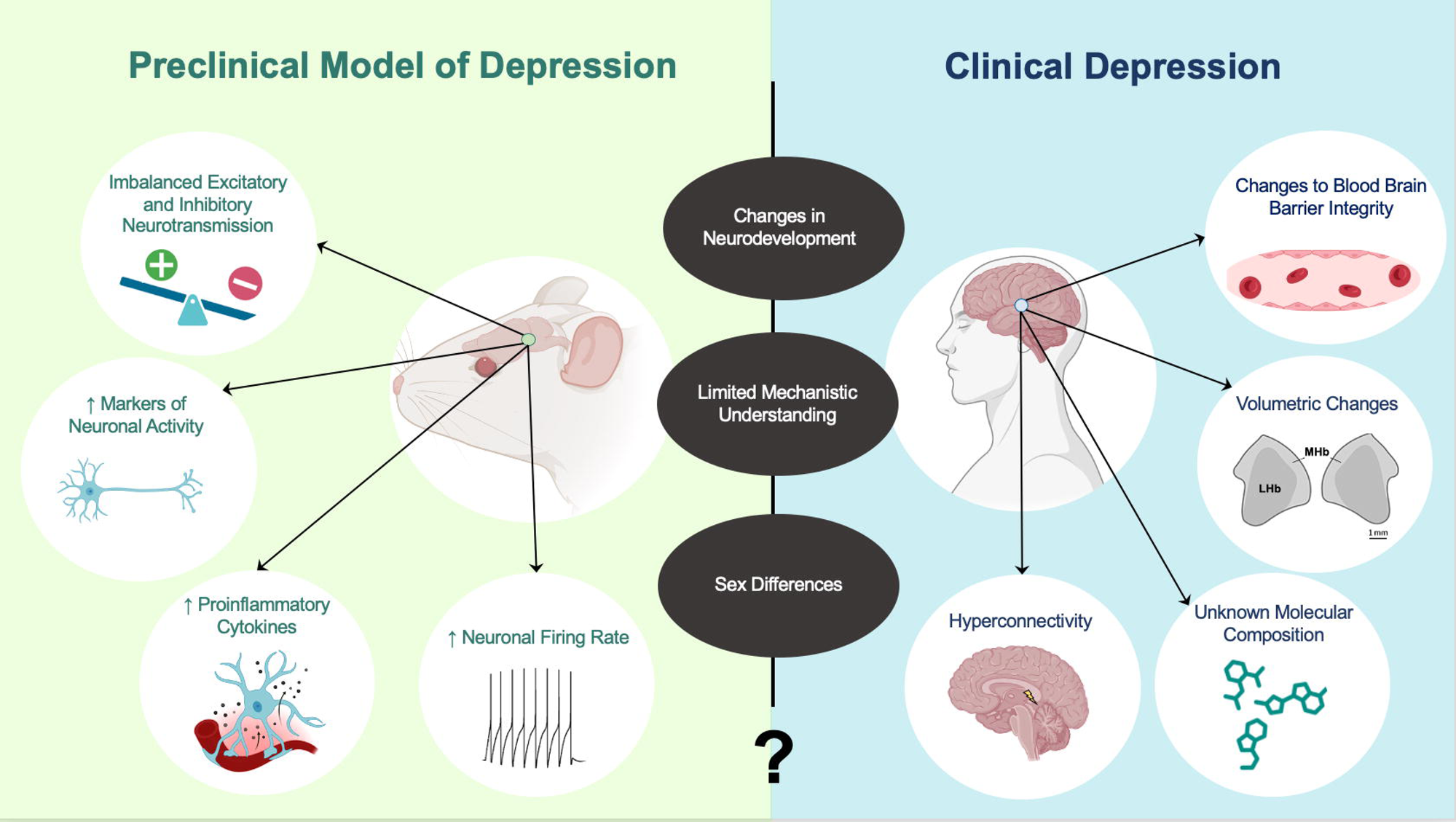

## Notes

### Competing Interest Statement

The authors have declared no competing interest.

